# Characterizing and minimizing the contribution of sensory inputs to TMS-evoked potentials

**DOI:** 10.1101/489864

**Authors:** Mana Biabani, Alex Fornito, Tuomas P. Mutanen, James Morrow, Nigel C. Rogasch

**Author notes:** Correspondence: Mana Biabani, Address: Monash University, Building 220, Clayton Campus, 770 Blackburn Rd. Clayton, VIC, 3800, Australia.

## Abstract

**Background:** Transcranial magnetic stimulation (TMS) evokes voltage deflections in electroencephalographic (EEG) recordings, known as TMS-evoked potentials (TEPs), which are increasingly used to study brain dynamics. However, the extent to which TEPs reflect activity directly evoked by magnetic rather than sensory stimulation is unclear.

**Objective:** To characterize and minimize the contribution of sensory inputs to TEPs.

**Methods:** Twenty-four healthy participants received TMS over the motor cortex using two different intensities (below and above cortical motor threshold) and waveforms (monophasic, biphasic). TMS was also applied over the shoulder as a multisensory control condition. Common sensory attenuation measures, including coil padding and noise masking, were adopted. We examined spatiotemporal relationships between the EEG responses to the scalp and shoulder stimulations at sensor and source levels. Furthermore, we compared three different filters (independent component analysis, signal-space projection with source informed reconstruction (SSP-SIR) and linear regression) designed to attenuate the impact of sensory inputs on TEPs.

**Results:** The responses to the scalp and shoulder stimulations were correlated in both temporal and spatial domains, especially after ∼60 ms, regardless of the intensity and stimuli waveform. Among the three filters, SSP-SIR showed the best trade-off between removing sensory-related signals while preserving data not related to the control condition.

**Conclusions:** The findings demonstrate that TEPs elicited by motor cortex TMS reflect a combination of transcranially and peripherally evoked brain responses despite adopting sensory attenuation methods during experiments, thereby highlighting the importance of adopting sensory control conditions in TMS-EEG studies. Offline filters may help to isolate the transcranial component of the TEP from its peripheral component, but only if these components express different spatiotemporal patterns. More realistic control conditions may help to improve the characterization and attenuation of sensory inputs to TEPs, especially in early responses.

## Introduction

The combination of transcranial magnetic stimulation (TMS) and electroencephalography (EEG) has become an increasingly popular technique, since it has extended the application of TMS to study brain dynamics across the cortex (Chang et al., 2018; Ferreri & Rossini, 2013). TMS-evoked potentials (TEPs) appear as voltage deflections in EEG recordings time-locked to the TMS pulse, which occur within a 300 to 500-ms long time window following stimulation, and can be used to make inferences about cortical reactivity and connectivity (Ilmoniemi et al., 1997). Recent studies have suggested that certain TEP peaks may reflect specific aspects of neurotransmission (Darmani et al., 2016; Manganotti, Acler, Masiero, & Del Felice, 2015; Premoli et al., 2014). For instance, pharmacological evidence has shown that earlier TEP peaks following stimulation of motor cortex (e.g. N45) are sensitive to neurotransmission mediated by GABA-A receptors, whereas later peaks (e.g. N100) are sensitive to GABA-B receptor mediated activity (Darmani et al., 2016; Nakazawa et al., 2012; Premoli et al., 2014; Treiman, 2001). However, the mechanisms underlying many aspects of TEPs are still not perfectly understood (Du et al., 2017; Rogasch & Fitzgerald, 2013). There is also debate over how much of the TEP signal represents direct activation of the cortex by TMS compared with other artefactual sources emanating from the recording equipment, blinks and muscle activity (Ilmoniemi et al., 2015; Ilmoniemi & Kičić, 2010; T. P. Mutanen et al., 2016; Rogasch et al., 2014).

One potential confound of TEPs is the interaction of TMS with the sensory system (Ilmoniemi et al., 2015; T. P. Mutanen et al., 2016; Rogasch & Fitzgerald, 2013; Rogasch et al., 2014). TMS is accompanied by a loud clicking noise, which results in auditory evoked potentials with peaks at similar intervals to TEPs, especially around N100 and P180. In addition, TMS activates sensory afferents in the underlying skin both mechanically by coil vibrations (e.g. a tapping sensation), and electrically by depolarizing the afferents fibers of cranial and facial nerves, which results in somatosensory-evoked potentials (Ilmoniemi & Kičić, 2010; Paus, Sipila, & Strafella, 2001). These peripherally-evoked potentials (PEPs) are often minimized during the experiment by playing white noise to mask the TMS click, and by using a layer of foam between the coil and scalp to dampen coil vibration (Ilmoniemi & Kičić, 2010; Massimini et al., 2005). Although these methods are commonly assumed to render the distorting effects of PEPs negligible (Ilmoniemi & Kičić, 2010; Nikouline, Ruohonen, & Ilmoniemi, 1999; Paus et al., 2001; Ter Braack, de Vos, & van Putten, 2015), several recent studies have demonstrated similarities between TEPs following TMS and realistic control conditions (e.g. stimulation of the shoulder or electrical stimulation of scalp with a TMS click) despite the PEP-masking procedures (Conde et al., 2018; Gordon, Desideri, Belardinelli, Zrenner, & Ziemann, 2018; Herring, Thut, Jensen, & Bergmann, 2015). These findings raise concerns regarding the specificity of TEPs to TMS-evoked cortical activity and underscore the urgent need for methods to further suppress sensory-evoked activity in TEP recordings.

There were two aims of the current study. The first aim was to characterize the contribution of sensory inputs to TEPs following stimulation of motor cortex. The second aim was to assess different offline methods for suppressing sensory responses in motor TEP recordings. We compared the efficacy of three different filtering methods in suppressing PEPs, as they have shown success in suppressing other types of artefacts such as ocular, decay and muscle artefacts (Rogasch et al., 2014; Verleger, Kuniecki, Möller, Fritzmannova, & Siebner, 2009). Finally, we repeated the above procedures by changing the intensity and waveform of the stimulations to examine the generality of the effects across different stimulation parameters.

## Methods

### Participants

Two separate experiments were conducted for this study. A total of 24 right-handed healthy individuals between the ages of 18 and 40 years were recruited. Twenty participants took part in experiment I (24.50 ± 4.86 years; 14 females) and 16 participants (25.82 ± 5.99 years; 11 females) participated in experiment II. Twelve individuals were common between experiments. All participants were screened for any contraindications to TMS (Rossi, Hallett, Rossini, Pascual-Leone, & Group, 2009), and provided their written consent prior to testing. All procedures were approved by the Monash University Human Ethics Committee in accordance with the declaration of Helsinki. Participants were seated comfortably with their elbows resting on the armrests, and their forearms pronated and rested on a pillow on their lap. They were also asked to keep their eyes open and focus on a black screen in front of them.

### EMG

Electromyographic (EMG) activity was recorded from the right first dorsal interosseous (FDI) muscle, using bipolar surface Ag-AgCl electrodes (4-mm active diameter), placed in a belly-tendon montage with a distance of ∼2 cm. The ground electrode was positioned on the dorsum of the right hand, over the midpoint of the third metacarpal bone. EMG signals were amplified (x1000), band-pass filtered (10-1000 Hz), digitized at 5 kHz, epoched around the TMS pulse (−200 to 500ms).

### EEG

EEG recordings were made using a SynAmps^2^ EEG system (Neuroscan, Compumedics, Australia), from 62 TMS-compatible Ag/AgCl-sintered ring electrodes, embedded in an elastic cap (EASYCAP, Germany). The electrodes were positioned according to the 10–20 international system, online-referenced to FCz and grounded to AFz. Electrode positions were co-registered to each subject’s MRI by means of a neuronavigation system (Brainsight™ 2, Rogue Research Inc., Canada) and digitized. EEG signals were amplified (x1000), low pass filtered (DC – 2000 Hz), digitized at 10 kHz and recorded on a computer using the Curry8 software (Neuroscan, Compumedics, Australia), for offline analysis. The skin-electrode impedance level was maintained at below 5 kΩ throughout the session (Ilmoniemi & Kičić, 2010).

### TMS

In experiment I, biphasic TMS pulses were applied to the hand area of left M1, where stimulation could induce consistent MEPs with greatest amplitude in FDI muscle (Rossini et al., 2015), using a figure-of-eight coil (C-B60) connected to a MagPro X100+Option stimulator (MagVenture, Denmark). The stimulator was set to deliver biphasic pulses with anterior-posterior and then posterior-anterior current direction in the underlying cortex. The neuronavigation system was used to guide TMS coil positioning and improve consistency across stimulation trials. Resting motor threshold (rMT) was determined as the minimum TMS intensity required to elicit MEPs >50 μV in at least 5 of 10 consecutive trials (with EEG cap on), and was expressed as a percentage of maximum stimulator output (% MSO)(Rothwell et al., 1999). Each participant received 100 TMS pulses at an intensity of 120% rMT. As a control condition, 100 additional TMS pulses (120% rMT) were administered to participants’ shoulder over the left acromioclavicular joint to mimic auditory and somatosensory sensations experienced during TMS. We changed the orientation and angle of the coil until the participants reported the same level of local sensations under the coil between the real TMS and control conditions. While this control condition does not perfectly match the scalp sensation of TMS, it does control for the general sensory experience of the stimulation (Herring et al., 2015).

For scalp stimulations, the TMS coil was held tangentially over the left side of the scalp with the handle pointing backward with an angle of 45° with the sagittal plane. For shoulder stimulations, we changed the orientation and angle of the coil until the participants reported the same level of local sensations under the coil between the real and control conditions. We decreased the intensity for two participants since they reported propagation of the pulses towards the arm and hand, which couldn’t be prevented by changing the coil orientation. One subject, however, did not feel a strong enough tapping sensation on the shoulder so we increased the intensity to 135% rMT. On average, the stimulation intensity used for control conditions was not significantly different from 120% rMT (p = 0.8) (Table S1).

To investigate the effect of the stimulation intensity, each individual also received 100 TMS pulses with the intensity of 80% rMT over the left M1, in the same session. In experiment II, the effect of the waveform was explored by applying supra-threshold monophasic pulses over the left M1 and shoulder. All of the participants reported that a small remnant of the acoustic clicks was still perceivable during the stimulation blocks, even when the volume was set to their upper threshold of comfort.

During both scalp and shoulder stimulation conditions, all of the currently advised measures to minimize multisensory inputs were implemented (Ter Braack et al., 2015), including attaching a thin layer of foam underneath the coil to minimize coil vibration and bone-conducted auditory activation, and playing white noise through inserted earphones to minimize air-conducted auditory activation. For each individual, the intensity of the white noise was increased until the click sound produced by the stimulations at 120% rMT was unperceivable or the sound pressure reached their upper limit of comfort. All of the participants reported that a small remnant of the acoustic clicks was still perceivable during the stimulation blocks, even when the volume was set to their upper threshold of comfort (see supplementary methods for more details).

### EEG analysis

Analysis of EEG recordings was performed using custom scripts on the MATLAB platform (R2016b, The Mathworks, USA), EEGLAB (Delorme & Makeig, 2004), and TESA (Rogasch et al., 2017) toolboxes. The pipeline for cleaning and analyzing TMS-EEG data was based on the method described in (Rogasch et al., 2017; Rogasch et al., 2014) (supplementary methods). Cortical sources of the evoked potentials were estimated using the Brainstorm (v3) software (Tadel, Baillet, Mosher, Pantazis, & Leahy, 2011) and customized MATLAB scripts employing both minimum norm estimation (MNE) and dipole fitting methods (detailed in supplementary methods). All code for EEG processing and statistical analyses is available at https://github.com/BMHLab/TEPs-PEPs, and all clean data can be downloaded from https://doi.org/10.26180/5c0c8bf85eb24. The raw data is also available upon request.

### Suppression of PEPs

In order to suppress the contribution of PEPs from TEPs, we applied three different filtering techniques to each individual’s responses: 1) linear regression; 2) independent component analysis (ICA); and 3) signal-space projection with source-informed reconstruction (SSP-SIR). Linear regression involved obtaining the line of the best fit between the PEPs (from the control condition) and TEPs (from the real condition) at each point of time across electrodes, followed by subtracting the fitted PEP curves from the TEP data. For ICA, TEPs and PEPs from each individual were concatenated and submitted to the FastICA algorithm (Hyvärinen & Oja, 2000) (in addition to the two ICA steps already made in EEG pre-processing). SSP-SIR is a spatial filtering method, originally designed to remove TMS-evoked muscle artefacts (T. P. Mutanen et al., 2016). We applied SSP-SIR to TEPs by following all the steps detailed in (T. P. Mutanen et al., 2016). However, instead of using the frequency properties of muscle artefacts to estimate the projection matrix, we used the PEP data to estimate the artifactual dimensions to be removed from TEPs. To increase the accuracy, we applied the defined projection matrix to the lead-field matrix to take into account the distortions of the data in the inverse solution. For both ICA and SSP-SIR, we retained the *k* components that explained more than 90% of variance in PEP trials and removed those from the TEP data. In total, 8.6 ± 4.17 and 12.3 ± 2.34 components were rejected by ICA and SSP-SIR, respectively. (see supplementary for the topography of the rejected components by SSP-SIR).

A major issue in evaluating the success of artifact suppression methods is the lack of a ground truth (i.e. knowing what the TMS-evoked cortical activity should look like without PEP contamination). Therefore, we assessed the PEP suppression methods based on the assumption that the level of suppression should be related to the level of contamination (i.e. an ideal PEP suppression method should cause minimal distortion to TEPs when the relationship with PEPs is weak and vice versa). To test this assumption, we qualitatively compared TEPs before and after applying each filter at both scalp and source (estimations obtained using MNE) levels, and also measured their correlations across three different time windows. As a further assessment, we examined whether the quality of the source localization was altered using different filtering methods. Based on the assumption that short latency TEPs reflect localized activities around the site of stimulation, we expected that reducing contamination would improve dipole fitting at the earliest time point (N20). We assessed the quality of the dipole fit using a goodness of fit (GOF) measure (supplementary methods), and also compared the distance between the best-fit dipole source found in pre- and post-suppressed data at N20.

### Statistical analysis

All statistical analyses were performed in MATLAB. The signal-to-noise (SNR) of EEG recordings was estimated by dividing the maximum amplitude at each timepoint by the standard deviation of the signals recorded 100 ms prior to TMS trigger (Debener et al., 2007; Hu, Mouraux, Hu, & Iannetti, 2010). To compare the absolute values of TEPs and PEPs voltage levels across time and space, we applied cluster-based permutation tests over time and electrodes, as implemented in the FieldTrip toolbox (Oostenveld, Fries, Maris, & Schoffelen, 2011). To explore the relationships between the cortical responses to M1 and shoulder stimulations, we applied Spearman rank correlation tests across space (i.e. across electrodes at each time point) and time (i.e. across time for each electrode). Seven frequently studied peaks including N20, P30, N45, P60, N100, P180 and N280 were selected to examine the spatial correlations for each individual, and also to explore inter-individual variability in the TEP/PEP relationship. The spatial correlations were also examined for all points of time by testing the 95% of confidence intervals of correlation values against zero. The temporal correlations were assessed for each individual and each channel at three different time intervals (early, middle and late post-stimulus). To account for inter-individual variability in temporal properties of EEG responses, instead of defining fixed time windows for all subjects, the intervals were individualized according to the appearance of TEP peaks (i.e. N20-P60, P60-P180, P180-N280). For group level analyses, the correlation results were transformed to z using Fisher’s transform and statistical significance was assessed by applying one-sample permutation tests to test the null hypothesis that the individuals’ z scores at each peak or time interval were equal to zero (Charter & Larsen, 1983; Dunn & Clark, 1969). The family-wise error rate due to multiple testings across time and channels was controlled for by adjusting the p-values using the t_max_ method proposed by Blair & Karniski (Blair & Karniski, 1993). The z scores were subsequently transformed back to the original scale for presentation. The results of source-localization (GOF-values and TMS-target-to-dipole-location distance) were compared between groups using paired-sample permutation tests with 10,000 shuffles.

## Results

### TEP-PEP comparisons

Fig. 1 illustrates the spatiotemporal distribution of grand-average TEPs and PEPs to suprathreshold stimulation. All of the canonical TEP peaks were visible following M1 stimulation, with mean amplitudes ranging from ∼-8 to +8μV (Fig. 1A). In contrast, PEPs showed smaller voltage deflections ranging from ∼-4 to +4μV, with clear peaks at N100, P180 and N280, and only small peaks within the first 50 ms (Fig. 1B). Cluster-based permutation tests confirmed that TEPs were larger in amplitude than PEPs across time (Fig. 1C).

**Figure 1:**
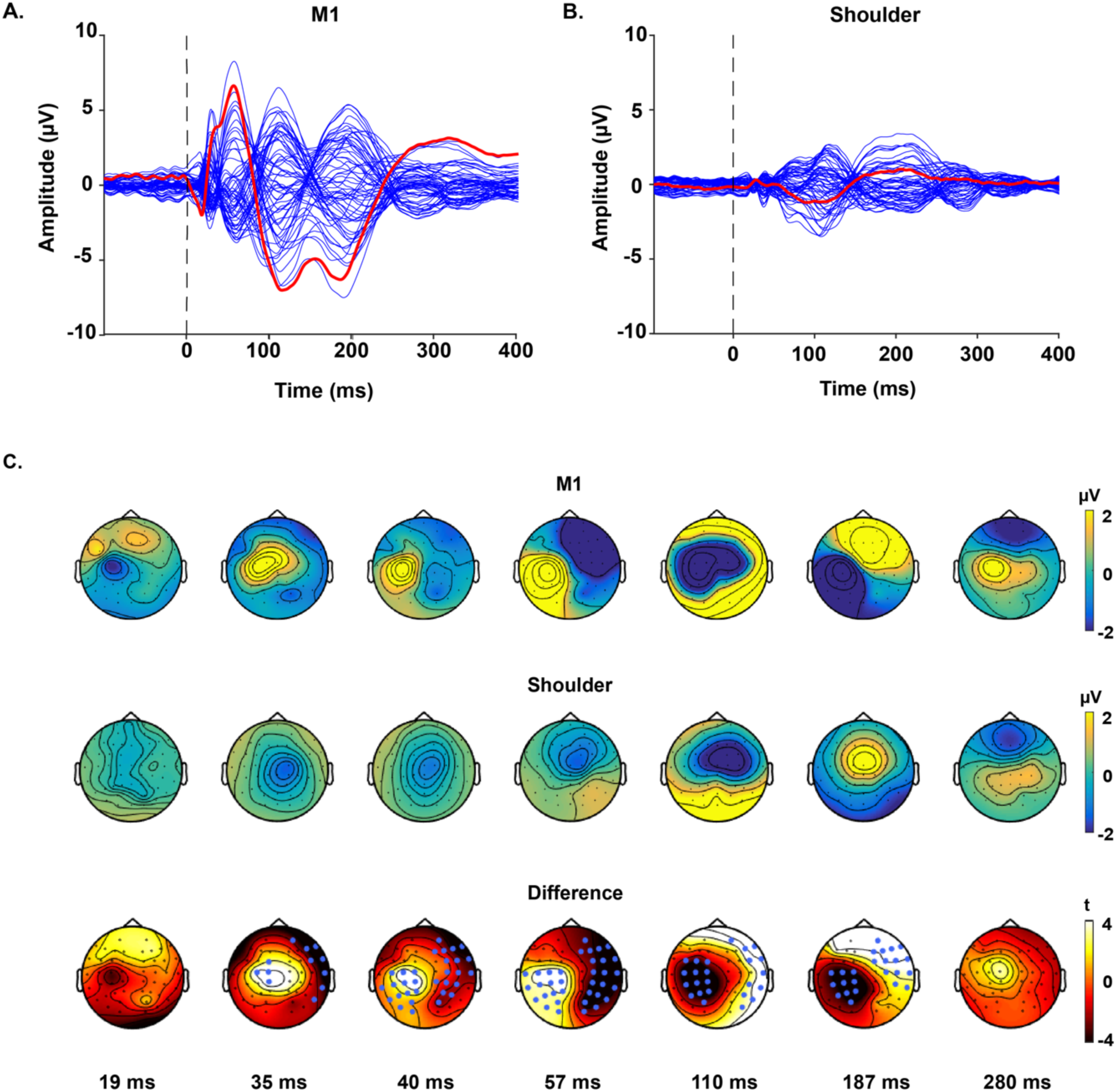
TMS-evoked potentials following suprathreshold, biphasic stimulation over left M1 and left shoulder. The butterfly plots show the grand-average of potentials recorded by each electrode. A) Responses to the stimulation of M1. B) Responses to the stimulation of shoulder. The red line indicates the recordings by the electrode underneath the coil (C3). The vertical dash line at zero time point indicates the point of time when TMS is applied. C) The upper and middle topoplots depict voltage distributions across the scalp for each peak of interest, in response to the real and control conditions, respectively. The lower topoplots illustrate the results of the cluster-based permutation tests comparing the voltage distribution of the two responses at each peak. Clusters were defined as at least two neighbouring electrodes exceeding the threshold of p-value < 0.05 at each point of time. Monte Carlo p-values were calculated on 5000 iterations with a critical α level set at p<0.025. The channels highlighted by blue dots belong to the clusters that showed statistically stronger responses to the real TMS condition. Two negative and three positive significant clusters were found.

Despite the amplitude differences between conditions, correlation analysis between spatial maps at each time point showed a consistent relationship (confidence intervals > 0) between TEPs and PEPs after ∼60 ms (Fig. 2A), suggesting a common underlying source after this time. We observed a positive correlation at baseline (pre-stimulus time window) between the two signals, which sharply dropped following the TMS trigger. Physiologically, we expected the correlations to fluctuate around zero (without offset) and then either remain around zero or increase in the presence of the common peripheral sensory inputs. To test whether the two-step ICA procedure adopted in our pre-processing pipeline could have caused a spurious correlation structure within the data, we repeated the analyses without ICA as performed in a previous study (Conde et al., 2019). As depicted in Fig. S1A, removing the ICA steps partially reduced the correlations during baseline and attenuated the sharp drop in correlation immediately following the TMS pulse. However, the general pattern of correlation between the signals was preserved, with consistent correlations between the signals present after ∼50 ms.

**Figure 2:**
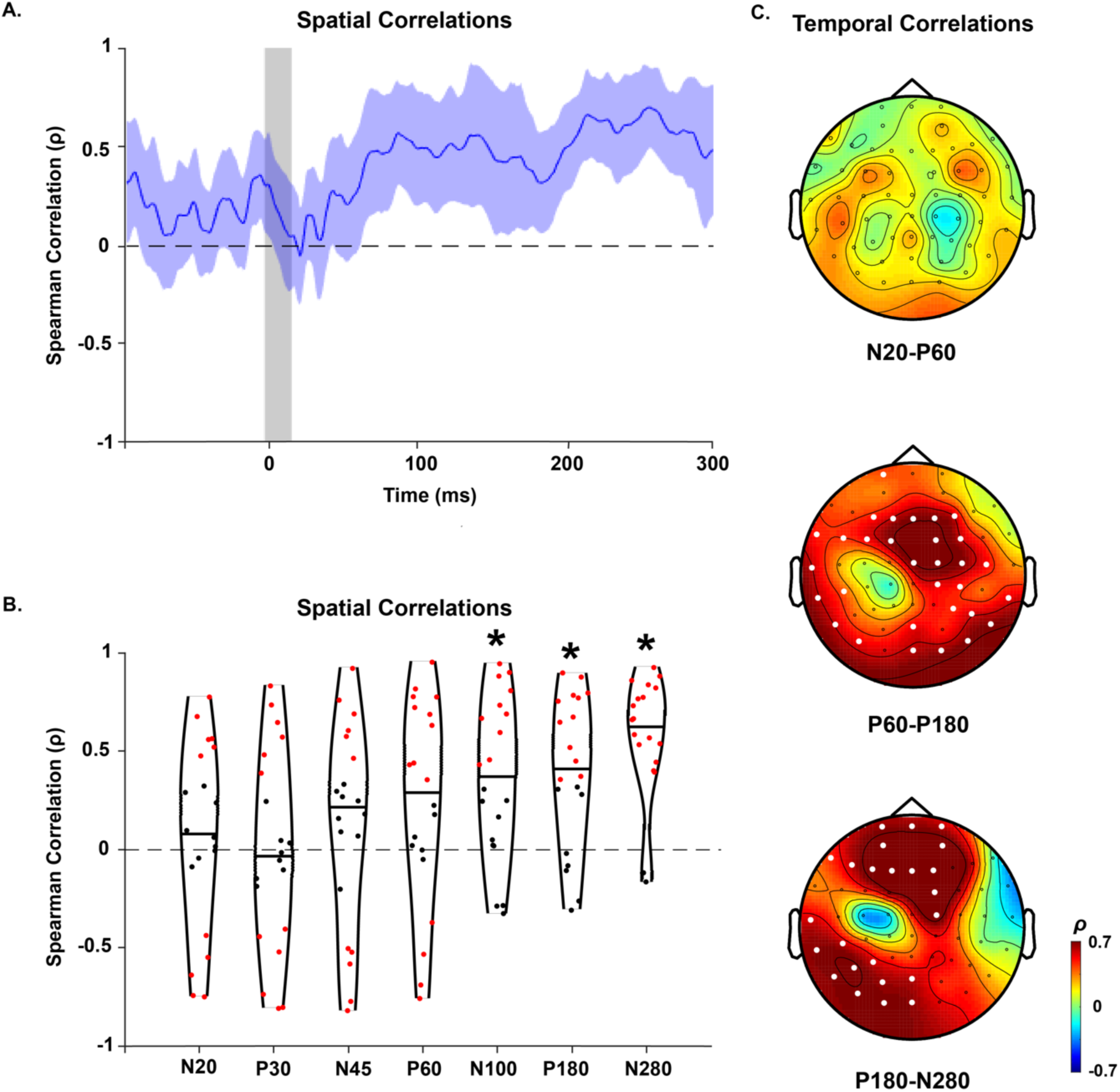
Spatiotemporal correlations of TEPs and SEPs. A) Spatial correlations between TEPs and SEPs at each point of time from 100 ms before to 300 ms following stimulations. The blue shaded area represents the 95% confidence intervals. The vertical grey bar shows the window of interpolated potentials around stimulus. B) The distribution of spatial correlations across individuals at the time of individualised TEP peaks. The dots within the violin plots represent the correlation values for each individual. The red dots above and below the zero line show significant positive and negative correlations, respectively (p<0.05), and the black dots represent non-significant correlations. * indicates that correlation values differed from 0 at the group level (one-sample t-test, p<0.05). C) The temporal correlations of the potentials at each window of time. White dots indicate the electrodes with significant positive correlations (p<0.05). No significant negative correlations were found.

As the exact timing of TEP peaks differ between individuals, we repeated the analysis using individualised peak times (Fig. 2B). The correlations between TEPs and PEPs showed high inter-individual variability at the earlier peaks, whereas the majority of participants showed significantly positive correlations between the conditions for the N100, P180 and N280 peaks. Analyses of the temporal correlations further confirmed the strong relationships between PEPs and TEPs after 60 ms. As depicted in Fig. 2C, none of the electrodes showed significant correlation between time series between 20-60 ms. However, a large number of electrodes showed positive correlations between 60-180 ms (35 of 62) and 180-280 ms (25 of 62). Of note, correlation values were highest over a cluster of fronto-central electrodes associated with PEPs. The lowest correlation values were found over a cluster centred over the stimulation site (M1). To further assess whether the motor cortex TMS condition contained signal with spatiotemporal patterns that differed from the shoulder stimulation condition, we applied principal component analysis (PCA) to the uncorrected data and compared the resulting components. The motor cortex condition contained a component which was not present in the shoulder stimulation condition and was strongest in electrodes over the site of stimulation and included peaks at ∼15, 60, 100 and 180 ms (Fig. 3).

**Figure 3:**
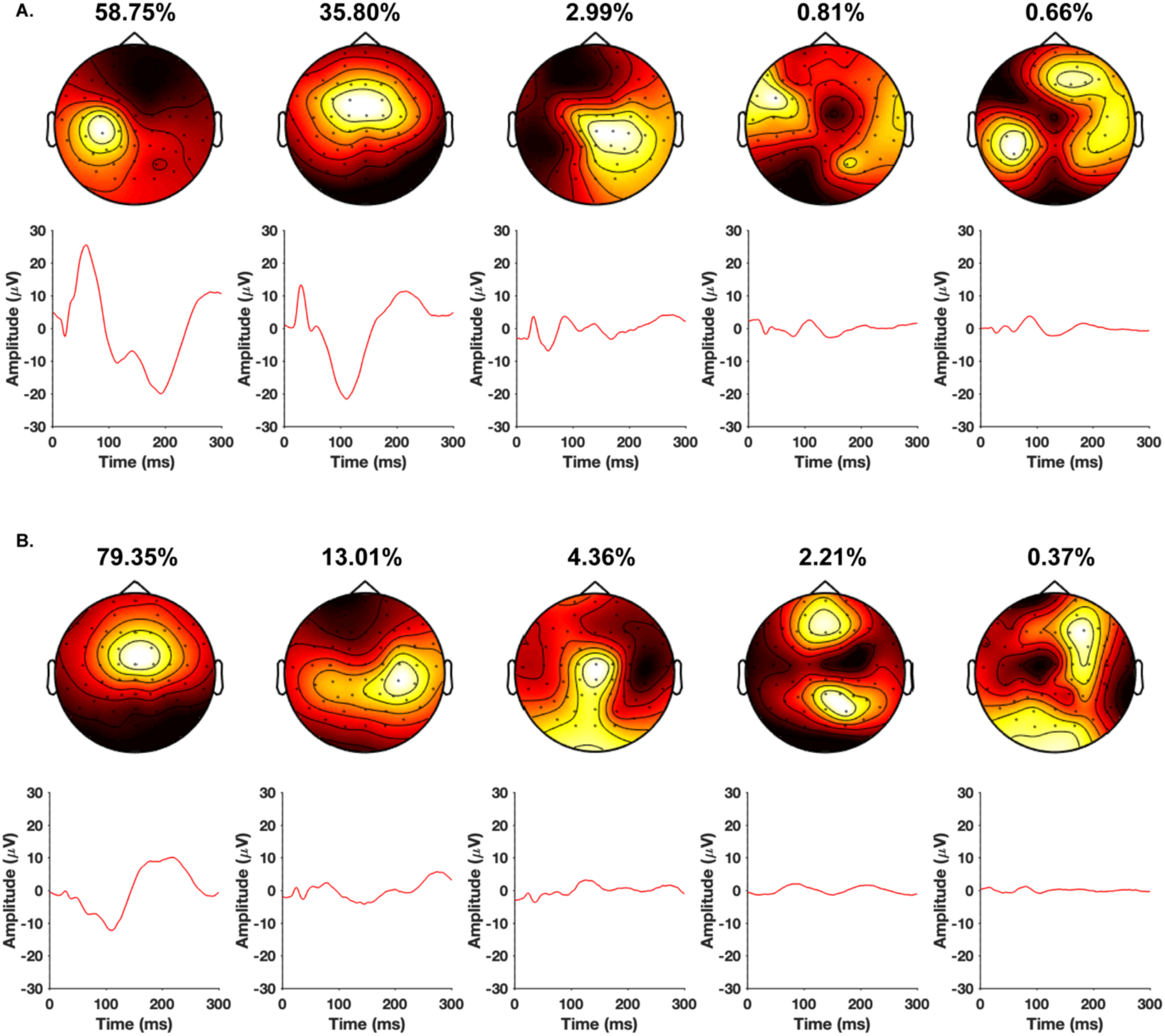
Principal components (explained 99% of the variance) of the signals evoked by suprathreshold biphasic stimulation. A) Stimulation is applied over M1 and B) Stimulation is applied over shoulder. The first component of M1 potentials, which explains the maximum variance of the signal (58.75%), cannot be found in control responses.

### Impacts of different PEP suppression methods on TEPs

Given that PEPs and TEPs showed high spatiotemporal correlation, we next compared three common methods for suppressing unwanted signals in TEP data: linear regression, ICA and SSP-SIR. Linear regression slightly diminished TEP amplitude, whereas, ICA exerted a strong impact on both temporal and spatial aspects of TEPs. The voltage range was considerably reduced and the peaks following 100 ms almost disappeared in C3 recordings. SSP-SIR preserved all the prominent peaks observed in the original data recorded by C3, but also caused a considerable reduction in voltage amplitude. While both linear regression and ICA reduced the SNR at almost all timepoints, SSP-SIR improved it at earlier peaks (Table 1). The spatial maps of the SSP-SIR-filtered TEPs demonstrated a fairly consistent pattern across all of the examined peaks centering the largest potentials near the site of stimulation (Fig. 4).

**Table 1:** Signal-to-noise ratio of each peak before and after each filtering method for suprathreshold stimulation condition (mean ± SD).

|  | **N15** | **P30** | **N45** | **P60** | **N100** | **P180** | **N280** |
| --- | --- | --- | --- | --- | --- | --- | --- |
| **Original** | 2.64<br>($\pm 2.61$) | 4.81<br>($\pm 6.84$) | 3.55<br>($\pm 3.72$) | 4.55<br>($\pm 3.17$) | 9.17<br>($\pm 7.34$) | 6.38<br>($\pm 3.85$) | 2.04<br>($\pm 1.83$) |
| **Linear**<br>**Regression** | 1.46<br>($\pm 1.13$) | 2.99<br>($\pm 2.93$) | 2.24<br>($\pm 1.99$) | 4.70<br>($\pm 4.39$) | 4.43<br>($\pm 4.04$) | 2.82<br>($\pm 2.87$) | 1.68<br>($\pm 1.06$) |
| **ICA** | 1.71<br>($\pm 1.15$) | 3.07<br>($\pm 3.12$) | 2.56<br>($\pm 2.30$) | 2.37<br>($\pm 1.84$) | 1.66<br>($\pm 1.29$) | 1.64<br>($\pm 1.76$) | 1.15<br>($\pm 1.05$) |
| **SSP-SIR** | 3.60<br>($\pm 4.40$) | 4.07<br>($\pm 4.43$) | 4.28<br>($\pm 4.65$) | 4.37<br>($\pm 3.40$) | 4.61<br>($\pm 6.61$) | 3.78<br>($\pm 3.01$) | 2.07<br>($\pm 1.59$) |

**Figure 4:**
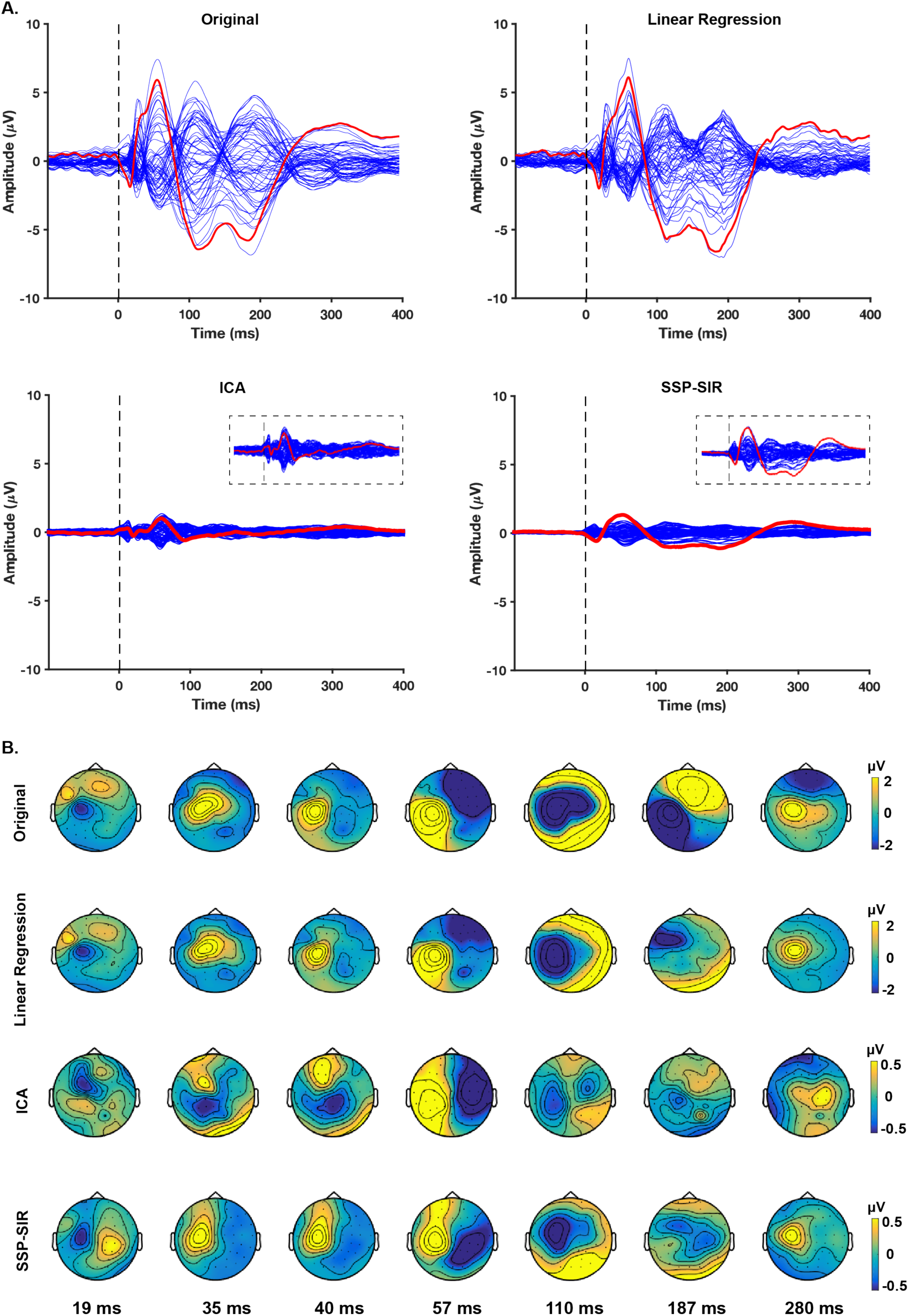
The alterations in the spatiotemporal distributions of TEPs induced by suprathreshold and biphasic TMS before and after removing PEPs using three different filtering methods. A) The butterfly plots demonstrate the grand-average of the potentials recorded by each electrode before (original) and after employing each filtering method. The red line indicates the recordings by the electrode underneath the coil (C3). The vertical dash line indicates the point of time when TMS is applied. Figure insets display a magnified view of the butterfly plots used to show the patterns in the small signals (i.e. ICA and SSP-SIR) more clearly (Y axis scale is set to [-2, 2]). B). The topoplots depict voltage distributions across the scalp for each peak of interest before (original) and after applying each filter. The colour bars have been re-scaled following ICA and SSP-SIR to indicate the spatial distribution of the small TEPs more clearly.

To compare the impact of the filters on distributed source estimation, we applied MNE to the responses before and after TEP cleaning. Before filtering, the real TMS condition showed a focal cortical activity around the site of stimulation at N20, following which the estimated sources gradually scattered and spread across hemispheres, showing a similar distribution to the sources estimated following shoulder stimulation. As depicted in Fig. 5 the three filtering methods changed the pattern of the estimated source differently, with SSP-SIR constraining source activity close to the site of stimulation.

**Figure 5:**
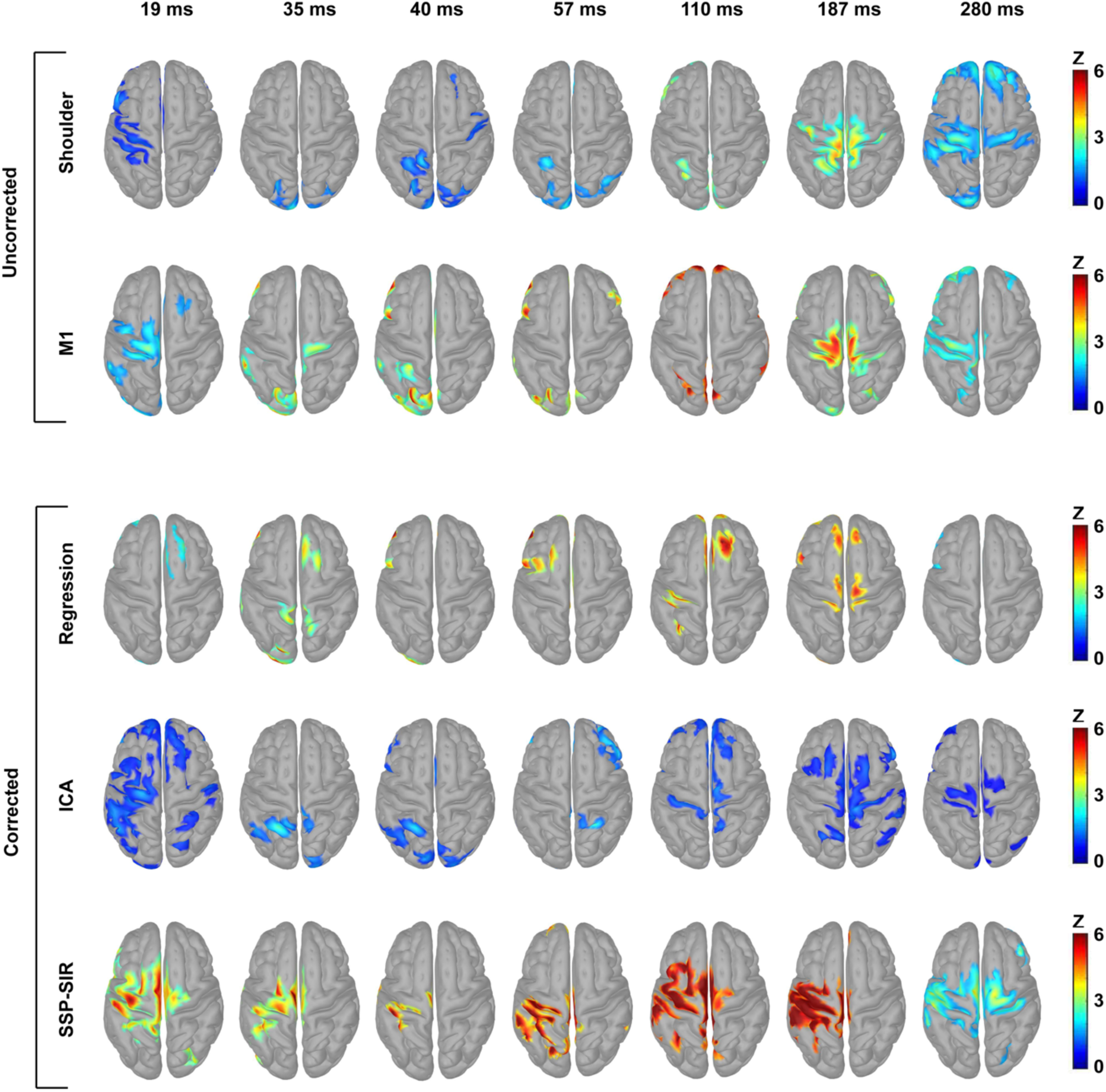
The estimated source distributions obtained by applying MNE to the TEPs from different filtering methods. MNE maps are thresholded at 40% of the maximum activity at each point of time and the minimum size for the activated regions is set to 50 vertices (for Linear Regression the active regions at P180 were smaller than 50 vertices; therefore, the minimum size was decreased to 30). Linear-regression altered the pattern of the estimated sources at some time points, importantly, at 20 ms when no change was expected. However, the pattern of source activation remained similar to shoulder stimulation for the later time points (e.g. N100 and P180). For ICA, sources are largely contained to left and right somatomotor cortices, sharing similarities with SEPs at some timepoints (e.g P180, N280). The estimated sources from SSP-SIR filtered TEPs are predominantly centered around the area under stimulation, substantially different from the pattern observed following shoulder stimulation.

Since using the same threshold level across conditions and/or time points could have caused non-optimal maps for visual comparisons, we carried out quantitative assessments of the filter-induced alterations to TEPs. We measured the correlation between TEPs before and after applying each filter at three different windows of time, at both sensor (for each channel) and source (for each vertex) levels (Fig. 6). At the scalp level, data following linear regression showed significant correlations with the original signal across all of time and space suggesting sub-optimal suppression of the sensory signal. A similar pattern was observed following ICA, although to a lesser extent at the later time window. SSP-SIR, however, resulted in low correlations (high suppression) especially around the fronto-central regions, which had shown high contamination. High correlations (low suppression) were observed at the recordings around the site of stimulation where TEPs had shown minimum correlations with PEPs. Similar to the scalp level, SSP-SIR showed low correlation with the original signal at the source level.

**Figure 6:**
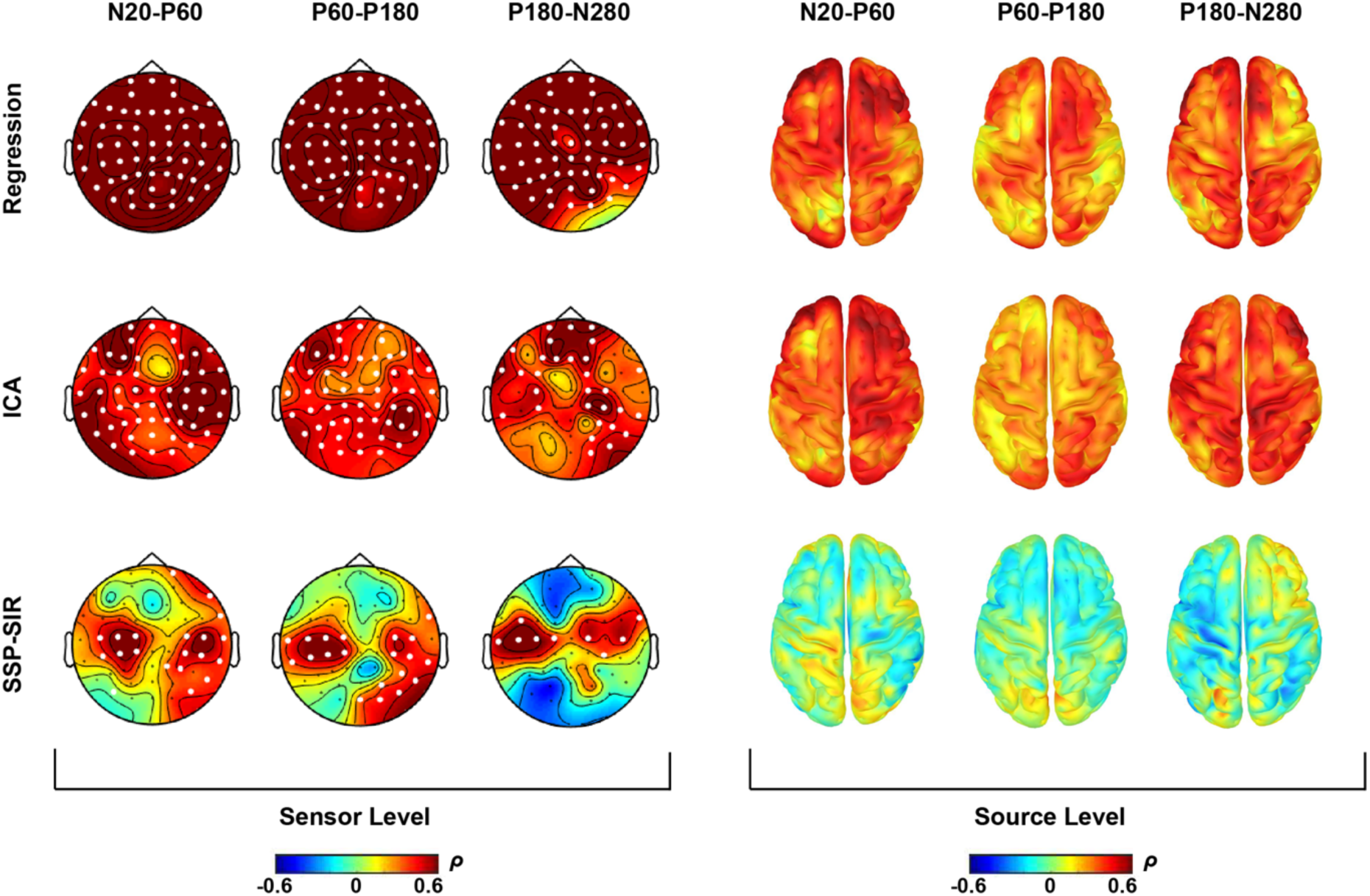
Spearman correlation measures between the original and filtered TEPs at both sensor and source levels at three different intervals. The maps show the average of the correlation values at individualized time windows (i.e. N20-P60, P60-P180, P180-N280). A) The correlations between the original and filtered potentials recorded by each channel at each window of time. White dots indicate the electrodes with significant positive correlations (p<0.05). B) The distribution of the correlations between the estimated source activities at each vertex.

To assess how each filtering method changed the quality of dipole fitting, we compared the best fit dipole found at N20 in the original and filtered data (Fig. 7). As depicted in Fig. 7, SSP-SIR substantially improved the GOF of TEP source estimations (permutation paired t-test with t_max_ correction, p>0.05), whereas the other filters reduced the GOF. All of the filtering methods resulted in significant displacement of the best fitting dipole (permutation paired-t test; p <0.05), while linear regression and ICA caused minimum and maximum repositions, respectively.

**Figure 7:**
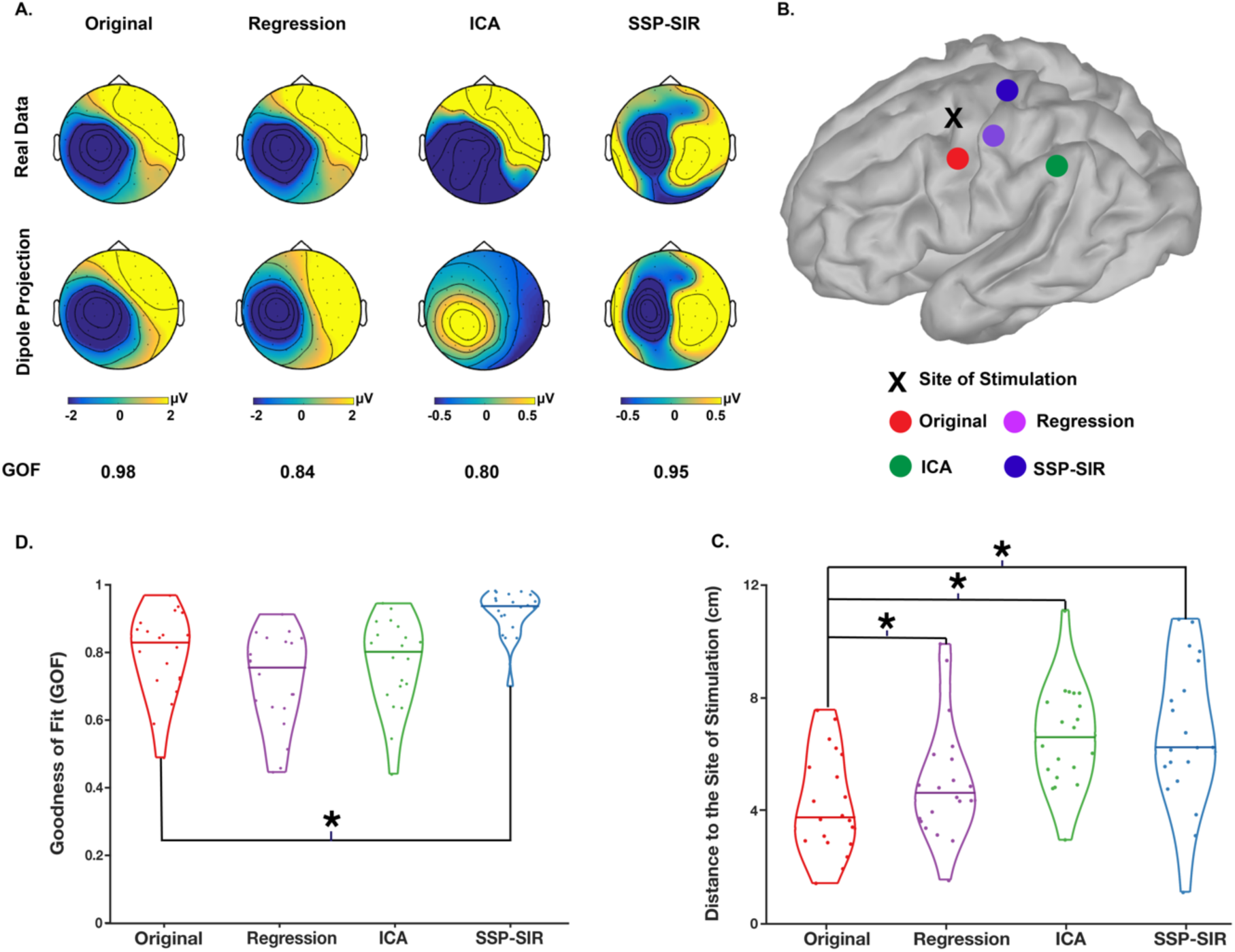
Comparisons of the effects of the different filtering methods on source localization at N20. A) Illustration of the measurements of goodness of fit (GOF), which represents the similarity between the topography of the real recorded data (upper plots) and the simulated responses produced by the dipole identified from each type of filtered data (lower plots), for one individual. B) Illustration of the measurements of the distances between the best-fit dipoles and the site of stimulation, for the same individual. C) Distribution of GOF for the best-fitting dipole source across individuals. * Indicates the significant change from raw data at the group level (P<0.05). D) Distance of the fitted dipole to the site of stimulation. The dots on violin plots represent the distance value for each individual. * Indicates the significant difference of the average distances between pre and post filtered potentials.

### Effect of stimulation parameters on TEP/PEP relationship and suppression

Decreasing the stimulation intensity substantially reduced the TEP amplitude relative to suprathreshold stimulation (to ∼±2μV) and changed the spatial distribution of voltages especially at the earlier peaks (i.e. Fig. 8). The spatiotemporal pattern of TEPs to subthreshold stimulation closely resembled PEPs to suprathreshold pulses, although the PEPs were significantly stronger at central electrodes, especially following ∼60 ms (Fig. 8).

**Figure 8:**
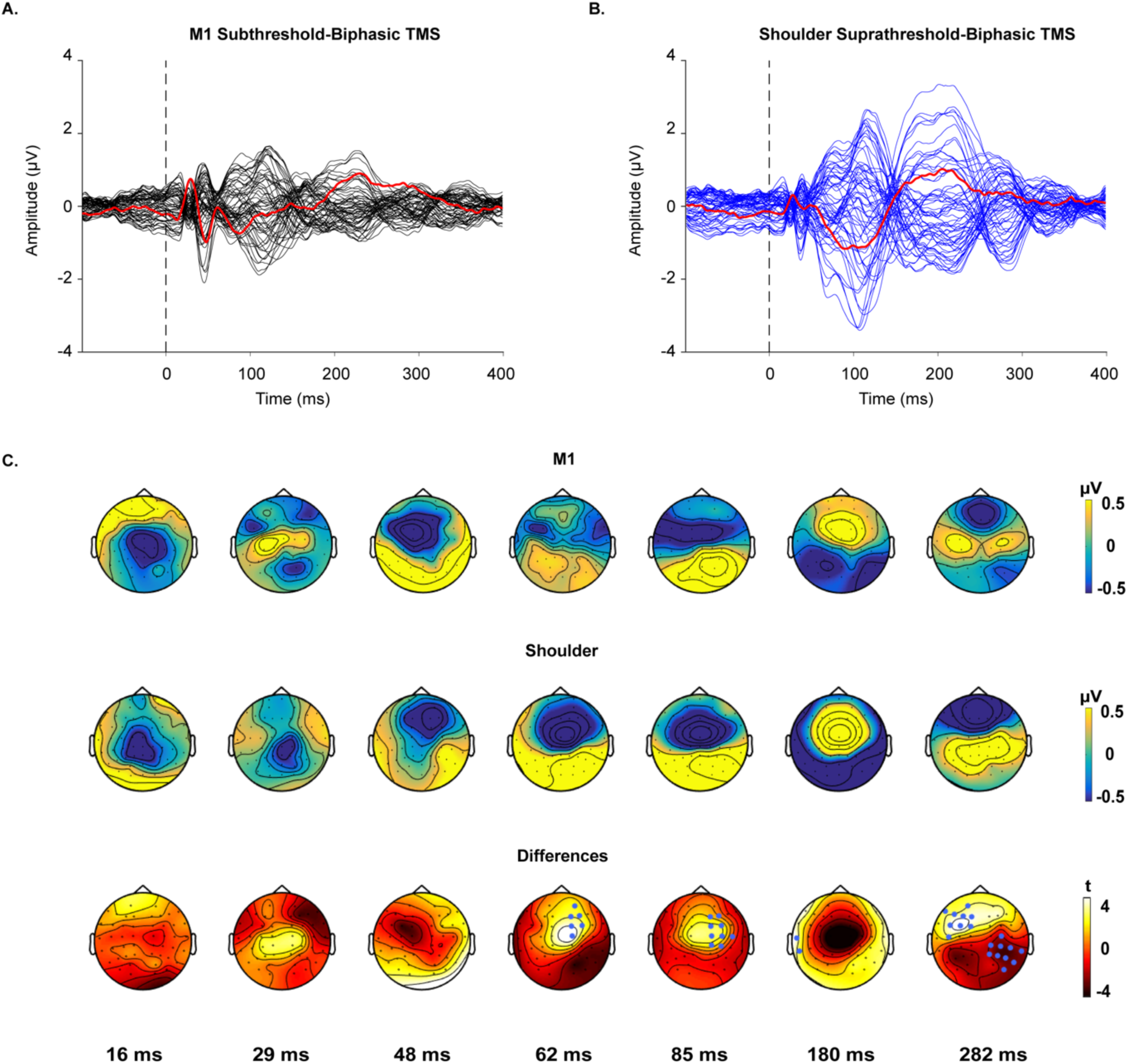
TMS-evoked potentials following subthreshold, biphasic stimulation over left M1 and suprathreshold, biphasic stimulation over left shoulder. The butterfly plots demonstrate the grand-average of potentials recorded by each electrode. A) Responses to the stimulation of M1. B) Responses to the stimulation of shoulder. The red lines indicate the recordings by the electrode underneath the coil (C3). The vertical dash line indicates the point of time when TMS is applied. C) The upper and middle topoplots depict voltage distributions across the scalp for each peak of interest, in response to the real and control conditions, respectively. The lower topoplots illustrate the results of the cluster-based permutation tests comparing the voltage distribution of the two responses at each peak. Clusters were defined as at least two neighbouring electrodes exceeding the threshold of p-value < 0.05 at each point of time. Monte Carlo p-values were calculated on 5000 iterations with a critical α level set at p<0.025. The channels highlighted by blue dots belong to the clusters that showed statistically stronger responses to shoulder stimulation (p<0.025). One negative and two positive significant clusters were found.

In addition, the two signals showed high spatiotemporal correlations after about 45 ms, with high inter-individual consistency after 60 ms (Fig. 9). The correlation at earlier timepoints improved after removing ICA from the pre-processing pipeline (Fig. S1B).

**Figure 9:**
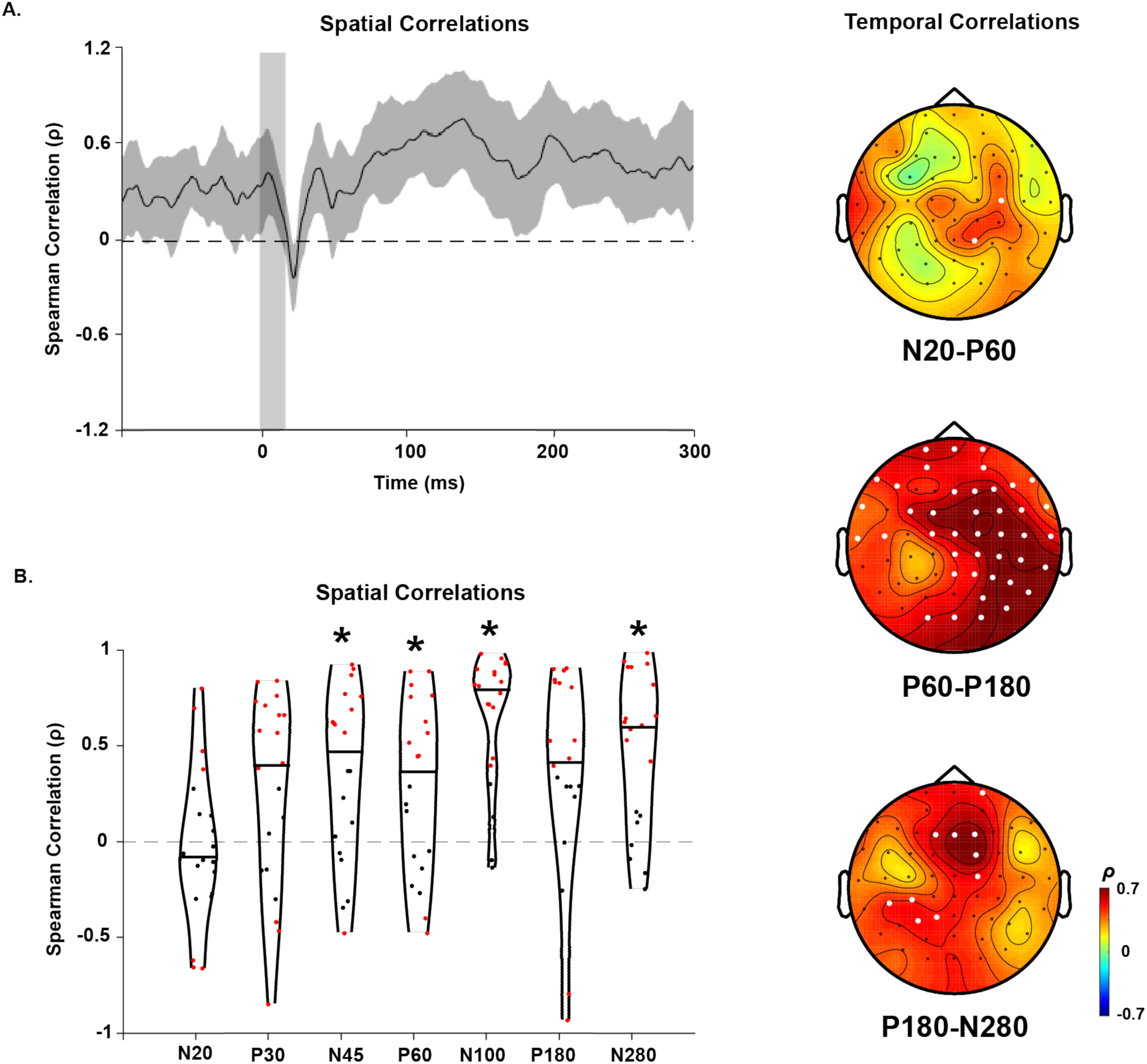
Spatiotemporal correlations of subthreshold TEPs and suprathreshold PEPs. A) The spatial correlations of the potentials at each point of time from 100 ms before to 300 ms following stimulations. The grey shaded area represents the 95% CIs. The vertical grey bar shows the window of interpolated potentials around stimulus. B) The distribution of spatial correlations across individuals. The dots within the violin plots represent the correlation values at for each individual. The red dots show significant positive and negative correlations respectively (p<0.05) and the black dots represent non-significant correlations. * indicates that correlation values differed from 0 at the group level (one-sample t-test, p<0.05). C) The temporal correlations of the potentials at each window of time. White dots indicate the electrodes with significant positive correlations (p<0.05). No significant negative correlation was found.

Despite the stronger responses to the control condition, SSP-SIR did not entirely suppress TEP peaks, especially at the earlier time points (<60ms) (Fig. 10). The global mean amplitude remained above baseline (baseline = 100 ms pre-stimulation, threshold = mean±3SD of baseline, P = 0.009) and the SNR improved at some early peaks (Table 2).

**Figure 10:**
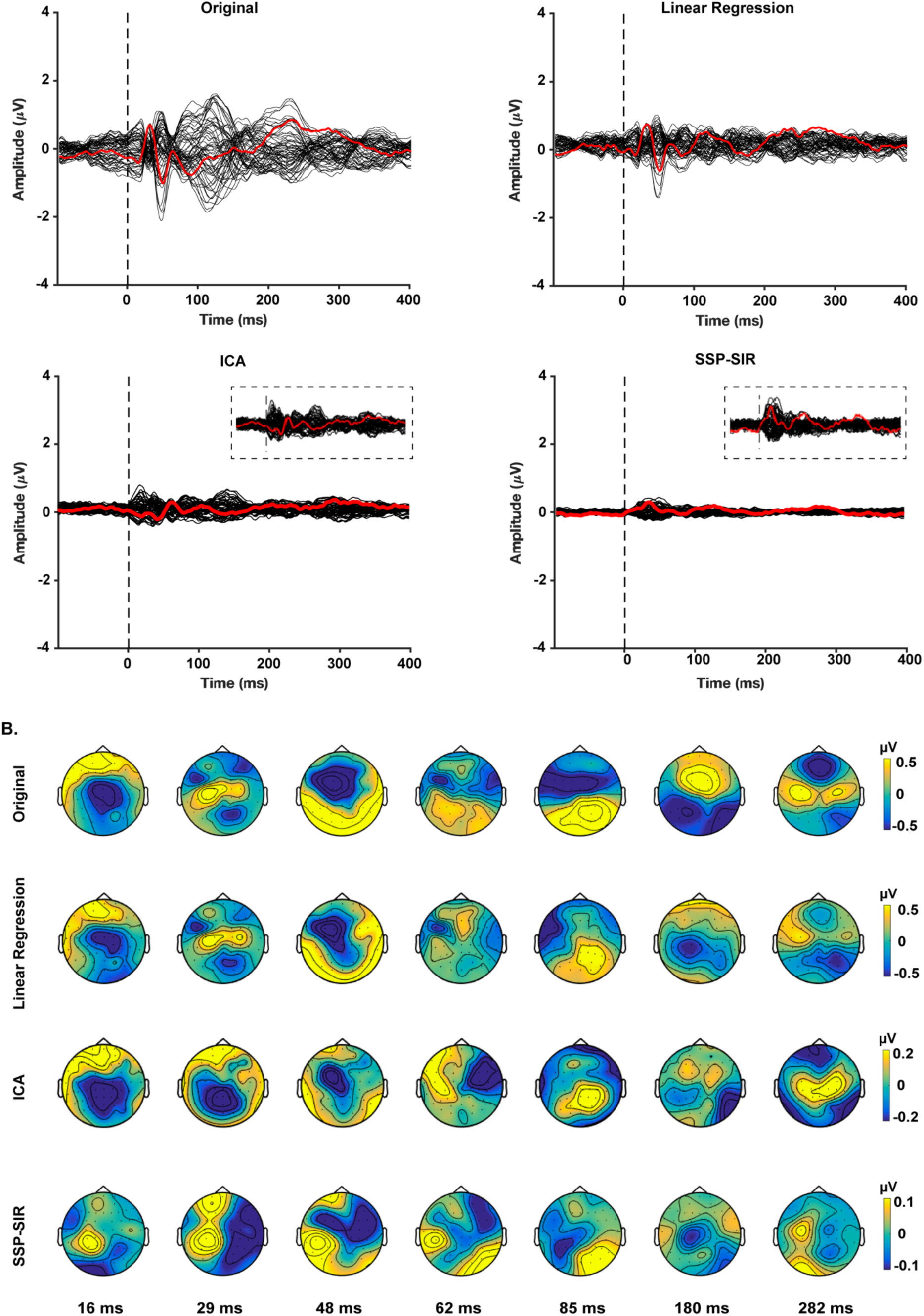
Alterations in the spatiotemporal distributions of TEPs induced by subthreshold and biphasic TMS before and after removing PEPs using three different filtering methods. A) The butterfly plots demonstrate the grand-average of the potentials recorded by each electrode before (original) and after employing each filtering method. The red line indicates the recordings by the electrode underneath the coil (C3). The vertical dash line indicates the point of time when TMS is applied. Figure insets display a magnified view of the butterfly plots used to show the patterns in the small signals (i.e. ICA and SSP-SIR) more clearly (Y axis scale is set to [-0.5, 0.5]). B). The topoplots depict voltage distributions across the scalp for each peak of interest before (original) and after applying each filter. The colour bars have been re-scaled following ICA and SSP-SIR to indicate the spatial distribution of the small TEPs more clearly.

**Table 2:**
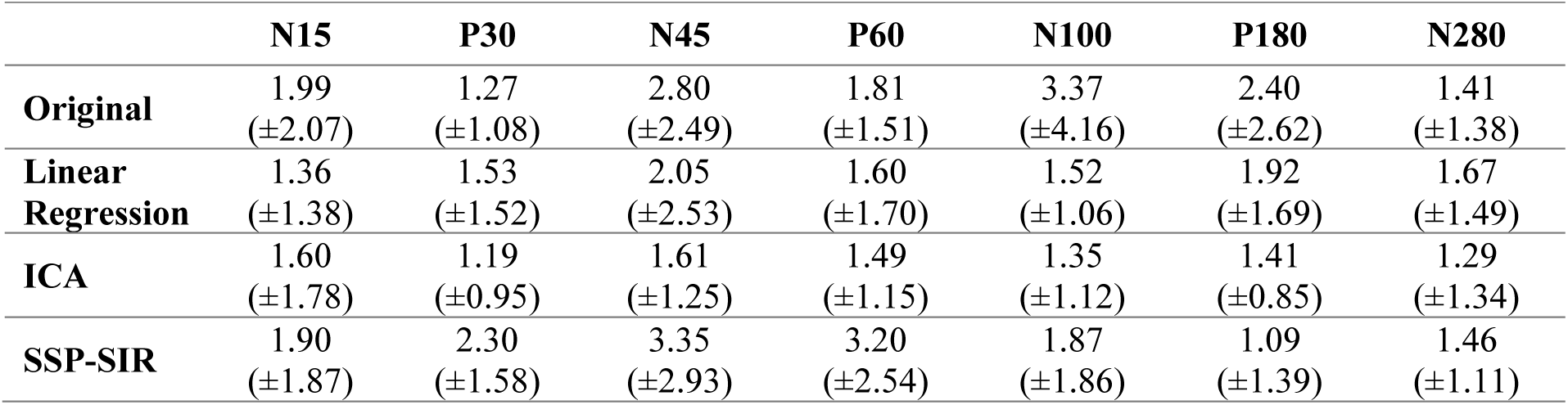
Signal-to-noise ratio of each peak before and after each filtering method for subthreshold stimulation condition (mean ± SD).

We also assessed how the filters impacted differences between suprathreshold and subthreshold TEPs. Linear regression did not alter the pattern of difference between TEPs from the two intensities, whereas, ICA and SSP-SIR substantially reduced the differences between stimulation intensities across time and space (Fig. 11 A). However, comparing the global mean field power (GMFP) between the two intensities indicated that suprathreshold TEPs were still larger in amplitude than subthreshold TEPs across time following all three filtering methods (p<0.05) (except for early responses following ICA which did not show significant differences) (Fig. 11 B).

**Figure 11:**
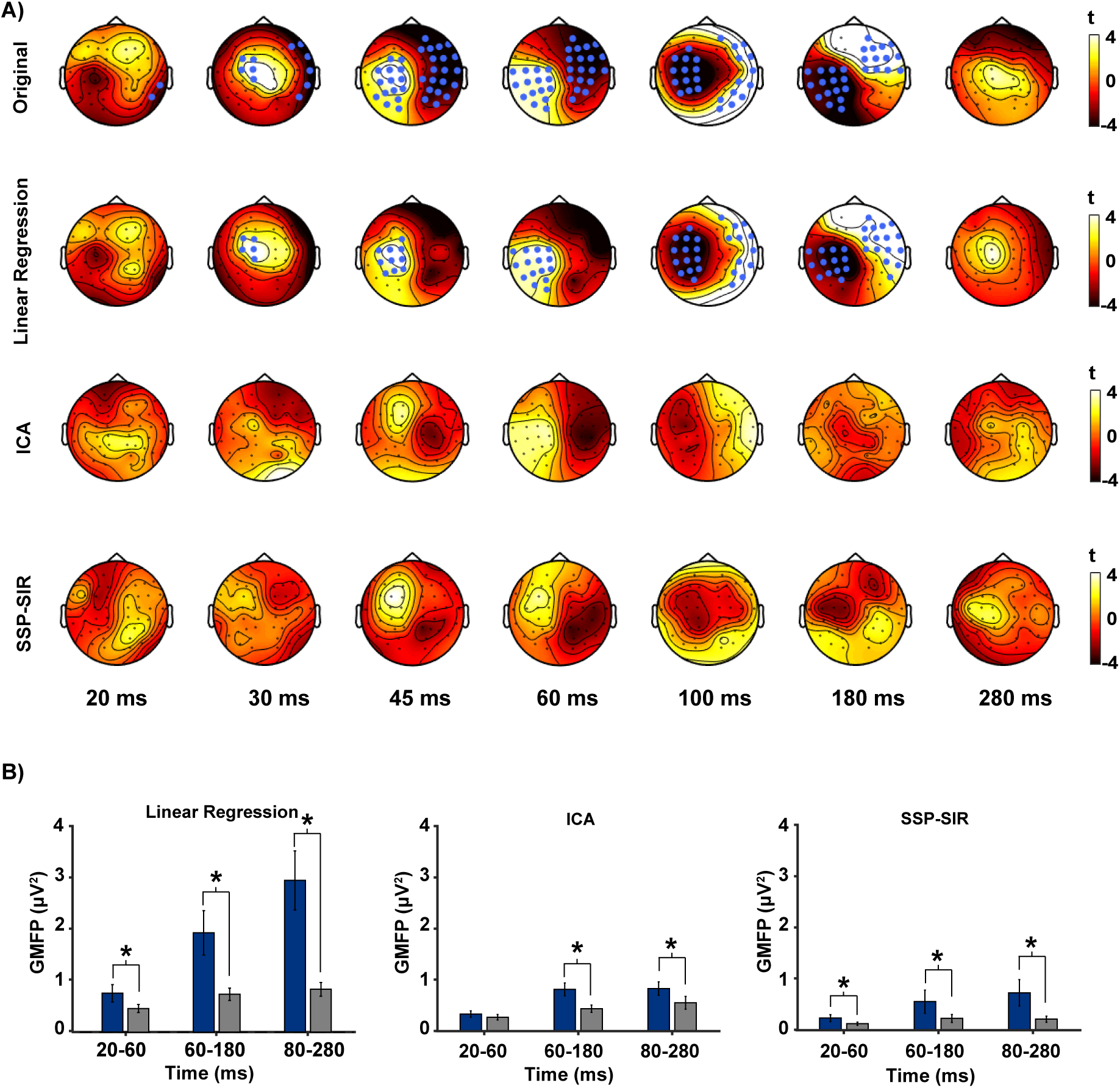
Comparisons of TEPs amplitude to sub- and suprathreshold TMS. A) The results of cluster-based permutation tests showing the spatiotemporal difference between the two responses before (original) and after sensory suppressions. The scalp maps illustrate the distribution of t values from cluster-based permutation tests at each evaluated peak. Clusters were defined as at least two neighbouring electrodes exceeding the threshold of p-value < 0.05. Monte Carlo p-values were calculated on 5000 iterations with a critical α level set at p<0.025. The channels highlighted by blue dots belong to the clusters that showed statistically stronger responses to the real TMS condition. Two negative and two positive significant clusters were found when comparing the original potentials. After suppressing PEPs using linear regression, one negative and two positive clusters were identified. No significant clusters were found between TEPs to sub- and suprathreshold TMS following ICA and SSP-SIR. B) Comparisons of GMFP of TEPs to sub- and suprathreshold TMS following the three filtering methods. * indicates significant difference between intensities (P<0.05).

Changing to the monophasic simulation waveform did not alter the spatiotemporal distribution of cortical responses compared to biphasic stimulation (n=12; P>0.025, cluster-based permutation tests). Similar to the suprathreshold biphasic condition, responses to monophasic stimulation over M1 were stronger than those to shoulder stimulation; however, at the alpha level of 0.025 the significant clusters were only found at N100 and P180 (Fig. S2). In addition, PEPs and TEPs evoked by monophasic TMS followed very similar correlation patterns as observed in the responses to biphasic stimulations, in both spatial and temporal domains (Fig. S3). SSP-SIR substantially decreased the potential amplitudes to ∼±1μV, preserving the original pattern of deflections and shifting the maximum amplitudes towards the site of stimulation at scalp and the majority of the peaks at source level, similar to the biphasic condition (Fig. S4-6).

## Discussion

In the present study, we investigated the contributing effect of sensory inputs to TMS-evoked potentials and investigated different offline filtering methods to suppress the impact of PEPs on TEPs. We found that, despite PEPs being lower in amplitude than TEPs induced by the same stimulation intensities, the two conditions were highly correlated in both space and time after ∼60 ms, suggesting that sensory input accounts for some of the spatiotemporal characteristics of TEPs at suprathreshold intensities. Among the three filtering methods tested to suppress PEPs in TEP recordings, SSP-SIR showed the highest efficiency in correcting the signals according to the pattern of contamination. Removing PEPs from TEPs revealed that even the late deflections in TEPs could not be fully explained by sensory inputs, suggesting TEPs at least partly reflect the direct cortical response to TMS. Changing the stimulation parameters (i.e. intensity and waveform) resulted in similar TEP-PEP correlations across time and space, and similar performance of PEP suppression methods. Our findings suggest that offline filtering methods such as SSP-SIR may be useful for attenuating PEPs in TEPs, but only if these components express different spatiotemporal patterns to avoid undesired suppression of the TEP of interest.

### Similarities and differences between TEPs and PEPs

In general, TEPs to subthreshold TMS showed higher spatiotemporal similarities with PEPs relative to suprathreshold stimulations across time. This may indicate a lower signal (real TEPs) to noise (PEPs) ratio following subthreshold stimulation, suggesting less sensitivity of EEG to detect TMS-evoked neural responses following subthreshold intensity TMS. TEPs from both intensities showed significant similarities with PEPs after 60 ms.

The N100 is perhaps the most extensively investigated peak in TMS-EEG recordings (Casula, Rocchi, Hannah, & Rothwell, 2018; Conde et al., 2018; Du et al., 2017; Johnson, Kundu, Casali, & Postle, 2012; Kaarre et al., 2018; Rogasch, Daskalakis, & Fitzgerald, 2015) and is thought to reflect cortical inhibition in both the motor and prefrontal cortices (Premoli et al., 2014; Rogasch et al., 2015). However, an N100 peak is frequently observed following a variety of sensory stimuli, including auditory (Scherg, Vajsar, & Picton, 1989; Ter Braack et al., 2015), somatosensory (Ter Braack et al., 2015; Wang, Mouraux, Liang, & Iannetti, 2008), visual (Mangun & Hillyard, 1991), pain (Greffrath, Baumgärtner, & Treede, 2007) and olfactory (Pause, Sojka, Krauel, & Ferstl, 1996) stimuli. Accordingly, an N100-P180 complex is observed following TMS pulses without cortical stimulation, resulting from the loud TMS click (air-conducted auditory component) and coil vibration (bone-conducted auditory and somatosensory component) (Nikouline et al., 1999; Nikulin, Kičić, Kähkönen, & Ilmoniemi, 2003; Tiitinen et al., 1999). As a result, it is common practice to apply noise masking through headphones and a layer of foam between the scalp and coil to mitigate these sensory inputs during TMS-EEG recordings (Ilmoniemi et al., 2015; Ter Braack et al., 2015). Despite applying these masking methods, we found considerable similarities between EEG responses to cortical and shoulder stimulation across different TMS intensities and pulse shapes. The observed relationship suggests that TEPs following motor cortex stimulation contain some potentials unrelated to the direct cortical responses to TMS, especially at later components (e.g. N100, P180 and N280).

While we observed a strong relationship between TEPs and PEPs after 60 ms, we did not observe a consistent relationship between the two signals across earlier time points (0-60 ms). Furthermore, TEPs from a cluster of electrodes over the site of stimulation did not correlate with PEPs across any time window. There are several possible explanations for this finding. First, the differences observed in the scalp pattern and amplitudes between cortical and shoulder responses could imply that TEPs following motor cortex TMS cannot be entirely explained by PEPs. If this explanation is correct, early TEPs (<60 ms) and TEPs from electrodes over the site of stimulation could reflect the true cortical response to TMS, which is not obscured by concurrent sensory responses. This explanation is supported by a recent study, which found differences between TEPs following subthreshold motor TMS and a realistic control condition (electrical scalp stimulation plus a coil click) when controlling for amplitude, particularly at early but not later peaks (Gordon et al., 2018). Whether the same is true for suprathreshold motor cortex stimulation requires further investigation.

Alternatively, the lack of relationship between TEPs and PEPs at early time points and in certain electrodes could result from discrepancies between other sensory inputs triggered by TMS, and which are not controlled by shoulder stimulation. For instance, TMS to motor cortex can result in face muscle twitches or stimulation of afferent nerves running across the scalp, subsequently leading to sensory responses in the face representation of the somatosensory cortex (T. Mutanen, Mäki, & Ilmoniemi, 2013). Moreover, motor-evoked potentials of the targeted muscle following suprathreshold pulses over M1 (right FDI here) can provide re-afferent sensory inputs to sensorimotor cortex (Fecchio et al., 2017). Following shoulder stimulation, it is likely that anti- and orthodromic spread of excitation along peripheral nerves causes muscle contractions and sensation in the upper limb. Due to the activation of different muscles with distinct spatial representations in cortex (Guyton & Hall, 2011) and the different conduction distance from face and limb muscles to the cortex, the somatosensory potentials induced by scalp and shoulder stimulations could have distinct spatiotemporal distributions, especially for early (<60 ms) responses. Sensory control conditions using electrical scalp stimulation may provide a more accurate representation of the somatosensory response to TMS than shoulder stimulation. Indeed, a recent study found higher correlation values between TEPs from both frontal and parietal cortex and PEPs at early time points (e.g. mean r values between ∼0.3- 0.55 at ∼45 ms following stimulation) than we have observed here. Whether this is due to differences in the stimulation site or control condition between studies remains unclear.

In addition to somatosensory differences, the level of auditory perception of the click differed between the scalp and shoulder stimulation. Stimulating the shoulder does not produce the same level of bone-conducted auditory input as that induced by scalp stimulation. Therefore, the amplitude of auditory-evoked potentials was likely smaller in the shoulder compared with the scalp TMS condition. However, it is worth mentioning that previous work using MEG suggests that differences in the sensory response to shoulder and face stimulation are subtle at the scalp level (Itomi, Kakigi, Hoshiyama, & Watanabe, 2001). Due to its inferior spatial resolution, EEG can be expected to be even less sensitive to this discrepancy. Moreover, the spatiotemporal pattern observed following shoulder stimulation in this study closely resembled the responses to the realistic control stimulations shown by previous studies (Conde et al., 2018; Gordon et al., 2018). Future work directly comparing different sensory control conditions both with each other (e.g. shoulder stimulation vs electrical scalp stimulation plus a coil click) and with TEPs following motor cortex stimulation is required to further disentangle the sensory contribution to motor TEPs.

Another important factor that may have altered the relationship between TEPs and PEPs in the earlier time window is the differences in cleaning procedures related to TMS-evoked muscle activity between the two signals. Activation of scalp muscles by TMS can result in high amplitude artefacts in the early EEG signal (<30 ms), which are often accompanied by decay artefacts related to electrode movement. These artefacts are not present following shoulder stimulation. We attenuated the muscle and decay artefacts in the TMS scalp condition using ICA. However, it is possible that ICA may also inadvertently remove some brain activity due to the temporal relationship between the signals (i.e. mixing of brain and artefact signal in the independent components removed), thereby altering the relationship between the real and control conditions artificially. Currently, it is difficult to assess the ability of ICA to accurately separate neural and non-neural signals. However, the development of more sophisticated and validated pre-processing methods for removing TMS-evoked muscle signals and related artefacts will help to rule out this limitation.

Taken together, our findings add to a growing body of evidence that a relevant part of the TEP response results from sensory input despite typical masking procedures. The relative contribution of the PEPs to the TEPs might have been underestimated in the present study due to differences in sensory inputs between scalp and shoulder stimulation, and different impacts of cleaning procedures on the two responses. More realistic control conditions (Conde et al., 2018) and more efficient pre-processing methods may improve the accuracy of comparisons between TEPs and PEPs, particularly in the earlier responses (<60 ms).

### Suppressing PEPs in TEP recordings

Given that online masking methods do not completely prevent contamination of TEPs by PEPs, we compared three offline methods for suppressing PEP activity: ICA, linear regression, and SSP-SIR. A major limitation in comparing these methods is the lack of a ground truth to benchmark performance against. Therefore, we evaluated each method based on the ability to correct later peaks with the focus on fronto-central regions, which were heavily contaminated by PEPs, without altering early signals especially around the site of stimulation, which were not as contaminated. Linear regression did not sufficiently reduce sensory contamination at later time periods. ICA was found to be more aggressive, strongly dampening the potentials (Fig. 4) and, more importantly, distorting the short latency peaks recorded around the site of stimulation (Fig. 6) (Li & Principe, 2006). Despite causing a substantial amplitude decrease, SSP-SIR showed a good trade-off between correcting the late and highly contaminated fronto-central potentials (Fig. 4, 6), while preserving the early potentials recorded at the least affected regions (Fig. 6, 7). Importantly, source estimation analysis following SSP-SIR correction were largely confined to the site of stimulation (M1), and to cortical areas that are strongly connected to M1, including ipsilateral supplementary motor area, ipsilateral premotor cortex, and contralateral M1, showing minimal overlaps with regions activated by shoulder stimulation (Fig. 5).

Separating two signals which are time-locked to the same event is an extremely challenging task, which relies on several assumptions. For instance, ICA assumes temporal independences of the underlying sources, an assumption which may not hold for time-locked signals such as TEPs and could explain why ICA removed so much of the signal. SSP-SIR also substantially attenuated TEP amplitudes before and after the TMS pulse at the scalp level. The reason for this strong attenuation is unclear but could either reflect that sensory signals account for a large amount of the overall EEG signal, or that PEPs and the genuine TEPs substantially overlap in space, leading to an undesired suppression of the TEPs of interest. The extent of this spatial overlap is likely to depend heavily on the stimulation site. Here we stimulated M1, which is located in the immediate vicinity of the primary sensory cortex, increasing the likelihood of overcorrection by spatial filters. However, PCA on the uncorrected data showed that there is a component in TEPs that is spatiotemporally independent from the control condition (Fig. 3), and therefore meets the requirements for using correction methods like SSP-SIR. Importantly, although the overall amplitude of the signal was attenuated, the global mean amplitude remained above baseline for both sub- and supra-threshold signals. Also, the SNR showed improvement following SSP-SIR, at both scalp (Table 1 and 2) and source levels (increased peak z scores (Fig. 5)). Whether this remaining signal reflects the true cortical response to TMS, or residual sensory activity not reflected in the shoulder control condition, such as re-afferent sensory input following MEP-related hand movement, requires further investigation.

Another key assumption which underlies all three suppression methods is the assumption of linear superposition of the sensory-evoked and TMS-evoked cortical activity. For this assumption to hold, the networks activated by sensory and TMS input would need to be completely independent, which is difficult to test quantitatively. As such, the suppression methods tested in this paper may not be appropriate for stimulation sites which overlap with sensory networks. Furthermore, to optimize sensory suppression with filtering methods, control conditions need to reproduce the PEPs resulting from scalp TMS as closely as possible. For instance, the accuracy of SSP-SIR as a spatial filter depends on how well the spatial features of the control data match the artefacts of interest. The sensory profiles of shoulder and M1 stimulation are not identical, which might have caused suboptimal sensory suppression in the current study. Future work comparing the effects of different control conditions on the outcomes of spatial filters, such as SSP-SIR, will help to determine which control condition is optimal for suppressing sensory potentials in TEPs. Regardless, comparing well-designed control conditions to TEPs is important to disentangle general sensory effects from specific TMS-evoked transcranial responses in TMS-EEG experiments.

## Conclusion

In conclusion, we have shown similarities between the cortical responses to TMS over M1 and the shoulder, especially from ∼60 ms following stimulation. The results imply that current practices for minimising auditory and somatosensory inputs during TMS do not completely eliminate the contribution of sensory inputs to TEPs from motor cortex and highlight the need for control conditions to detect and minimize these signals. However, TEPs could not be entirely explained by PEPs, either at early or late time points, implying that TEPs do contain signals reflecting the direct cortical responses to stimulation. Offline methods for suppressing sensory-evoked activity show promise for isolating TMS-evoked neural activity and will likely form an important step in obtaining cleaner and more reliable TEPs. However, these methods require that the transcranial-evoked and peripheral components of the TEP express different spatiotemporal patterns to avoid undesired overcorrection of the TEP of interest. More realistic control conditions may help to improve characterization and attenuation of sensory inputs to TEPs using offline filters, especially for early responses.

## Acknowledgments

This research was supported by the National Health and Research Council of Australia (<u>1072057</u>, NR; <u>1104580</u>, AF, NR) and the Australian Research Council (180100741, NR). The authors would like to thank Dr. Jaakko Nieminen and Dr. Ben D. Fulcher for valuable advice on data analysis.

