## Supplementary Materials for "Characterizing and minimizing the contribution of sensory inputs to TMS-evoked potentials"

### Supplementary methods

### TMS

In experiment II, monophasic TMS was delivered in the posterior-anterior direction through a figure-of-eight-shaped coil (7 cm wing diameter), connected to a Magstim 200 stimulator (Magstim Company, UK). This session was a part of a larger experiment in which the number of pulses was reduced (to an average of 75 pulses) to prevent the coil from overheating. Again, for this condition, the stimulation intensity used for control conditions was not significantly different from 120% rMT ($p = 0.11$) (Table S1). Since all participants had reported the perception of a click sound in the first experiment, we used a visual analogue scale (VAS) to quantify the efficiency of the white noise to cover the TMS click sound for this experiment. The VAS values ranged from 0 (could not hear the pulses at all) to 10 (the pulses were as loud as without protection), and the participants rated the sound level for each stimulation condition. The mean ± SD of VAS scores for suprathreshold M1 stimulations was 3.5 ± 1.21 and for shoulder stimulation was 1.81 ± 0.83 ($P<0.001$). None of the participants reported 0/10 click sound for any of the evaluated conditions. In all of the conditions, TMS pulses were given with the intervals that jittered between 4 and 6 s and the order of the real TMS and control conditions was pseudorandomized within each session.

### EEG analysis

First, the responses to the different stimulation conditions were concatenated together and epoched with a window of -1000 ms to 1000 ms around the TMS pulse, and the mean of each channel's baseline (defined as a window of -500 to -10ms) was removed from each epoch. The TMS pulse artifact (the data from -2 to 15ms) was then removed and a cubic interpolation was applied to replace the missing data. The recordings were then down sampled to 1000 Hz. Following which, the trials and channels affected by prominent artefacts were detected by visual inspections and removed (Table S1). Afterwards, the interpolated data around TMS pulse were replaced with constant amplitude data and the TMS induced muscle and decay artifacts were identified using the FastICA algorithm (Hyvärinen & Oja, 2000) and rejected. A linear interpolation was applied to replace the missing data. Data were then band-pass (1-100 Hz) and band-stop (48-52 Hz) filtered using a zero-phase Butterworth filter (order = 4) and other artifacts such as blinks and noise-related artefacts were corrected by applying a second run of FastICA algorithm. Artifactual components were selected using the TESA automated functions with default settings and checked visually (Table 1S). Finally, the rejected channels were spatially interpolated using spherical method and all data were re-referenced to the common average. TEPs and PEPs were computed for each participant and each condition by averaging the recordings over trials.

**Table S1** The average of rMT, stimulation intensities, and pre-processing outputs for each condition.

| | **Experiment I** | | | **Experiment II** | |
|---|---|---|---|---|---|
| | **120% rMT M1** | **80% rMT M1** | **120% rMT Shoulder** | **120% rMT M1** | **120% rMT Shoulder** |
| **rMT (mean ± SD)** | 56.8 (± 8.5) | 56.8 (± 8.5) | 56.8 (± 8.5) | 47.8 (± 6.6) | 47.8 (± 6.6) |
| **Intensity (mean ± SD)** | 68.2 (± 10.3) | 45.7 (± 6.9) | 67.9 (± 9.9) | 57.0 (± 7.9) | 55.6 (± 8.3) |
| **Remaining trials (mean ± SD)** | 78.0 (± 12.0) | 82.4 (± 9.5) | 81.4 (± 15.5) | 69.5 (± 23.5) | 70.2 (± 22.6) |
| **Remaining channels (mean ± SD)** | 61.6 (± 0.9) | 61.6 (± 0.9) | 61.6 (± 0.9) | 61.1 (± 1.2) | 61.1 (± 1.2) |
| **ICs removed (mean ± SD)** | 23.6 (± 4.9) | 23.6 (± 4.9) | 23.6 (± 4.9) | 23.3 (± 6.9) | 23.3 (± 6.9) |

rMT = Resting motor threshold; M1 = Primary motor cortex; ICs = Independent components

**Source estimation**

First, each individual's T1 scan was automatically segmented using FreeSurfer software (version 5.3). After visual inspections and manual corrections, the FreeSurfer output was imported to Brainstorm and the cortical surface was down sampled to 15,000 vertices. Registration between EEG and MRI was then performed by aligning the locations of EEG electrodes with the generated surfaces. Afterwards, the head model was computed using a three-layer symmetric Boundary Element Method (BEM; OpenMEEG freeware), applying the default conductivity values (i.e.  scalp = 1, skull= 0.0125 and brain= 1) (Gramfort, Papadopoulo, Olivi, & Clerc, 2010). The noise covariance matrix was then calculated following concatenation of the baseline periods (-1s to -0.002s pre-stimulus), for each trial, each individual and each condition separately. Afterwards, cortical sources were estimated using minimum norm estimation (MNE) (Depth weighting: 0.5, Regularize noise covariance: 0.1, SNR: 3), for which, sources were constrained to be normal to the surface of the cortex (Lin, Belliveau, Dale, & Hämäläinen, 2006). The source amplitude was then transformed into z-score relative to the baseline for each individual at each condition, and then projected to a common default anatomy (ICBM152) to facilitate averaging across participants and group analyses. The source values where then smoothed with the kernel size of 3mm for display purposes.

Source estimation was also performed using dipole fitting method for one selected point of time. For this method, 15000 freely oriented dipoles were positioned on the cortical surface. Dipole fitting was performed by minimizing the sum of squared errors between the scalp measured and projected data, and the dipole with the highest goodness of fit (GOF; i.e. the least error) was defined as the most likely location of the cortical source (Kaukoranta, Hämäläinen,

Sarvas, & Hari, 1986). The dipole fit was considered reliable if GOF value was over 0.90 (Mutanen, Metsomaa, Liljander, & Ilmoniemi, 2018). The individuals' selected dipoles were then projected to the default anatomy for group analyses.

### Supplementary results

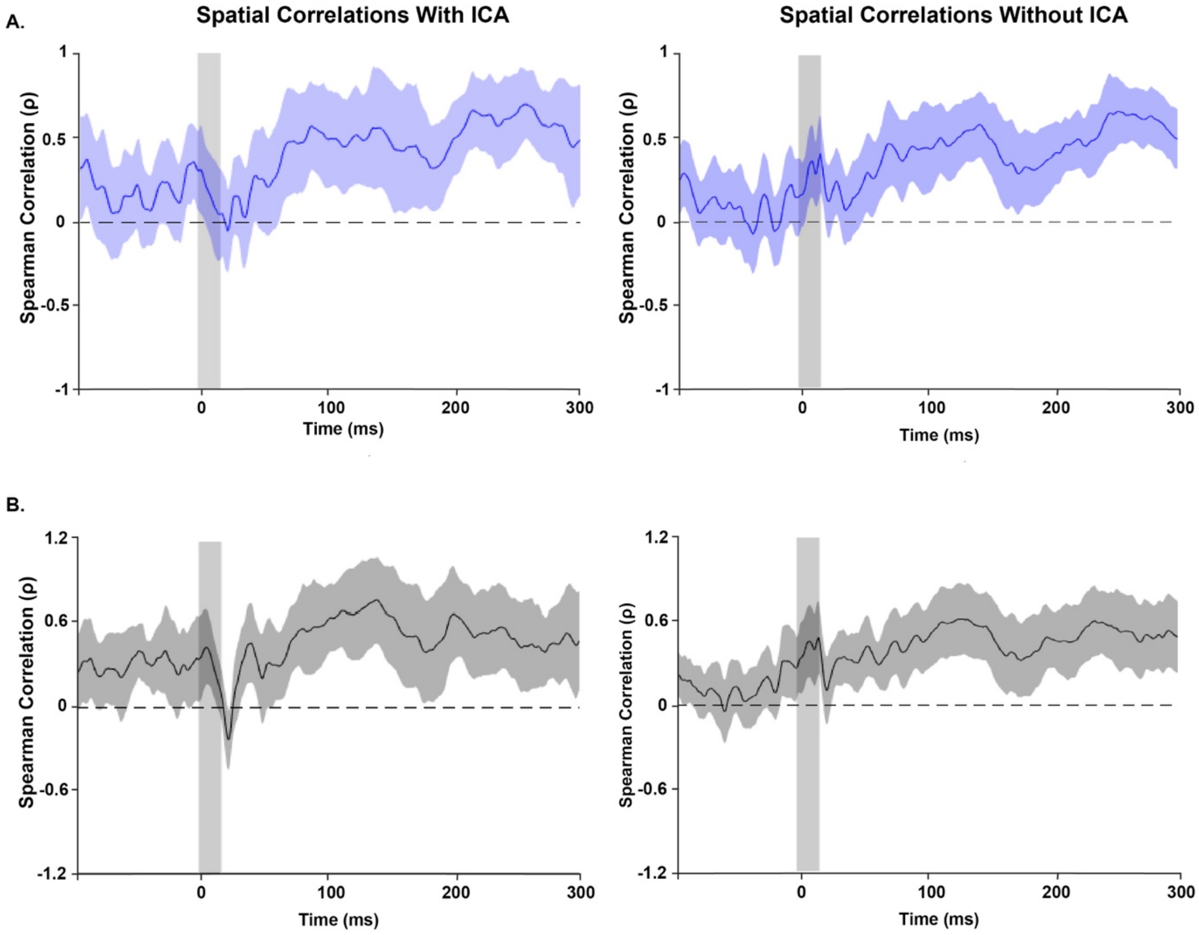


**Figure S1: Effect of excluding ICA from pre-processing pipeline on the spatial correlation between TEPs and PEPs.** A) suprathreshold and biphasic stimulation B) subthreshold and biphasic stimulation.

**Effect of stimulation parameters**

In addition to the assessments of different intensities, we repeated the experiment using Monophasic pulses to examine whether the observed TEPs-PEPs relationship generalises between waveforms. For which, suprathreshold monophasic pulses were employed for both real (stimulation over left m1) and control (stimulation over left shoulder) conditions. All of the assessments performed for biphasic stimulations were repeated for this condition.

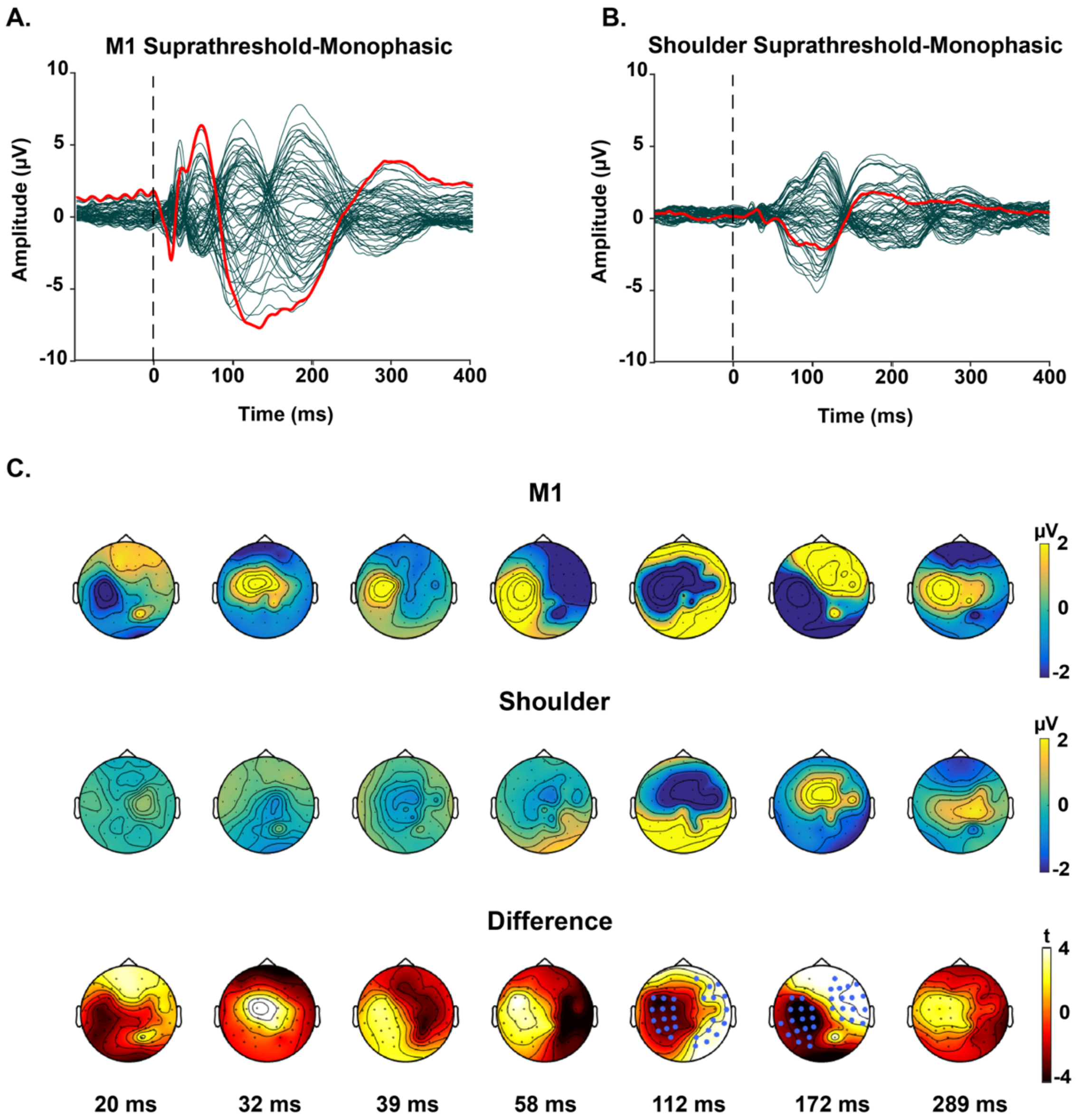

A.
M1 Suprathreshold-Monophasic
Amplitude (µV)
Time (ms)
B.
Shoulder Suprathreshold-Monophasic
Amplitude (µV)
Time (ms)
C.
M1
Shoulder
Difference
20 ms
32 ms
39 ms
58 ms
112 ms
172 ms
289 ms

**Figure S2: TMS-evoked potentials following suprathreshold, monophasic stimulations over left M1 and left shoulder.** The butterfly plots demonstrate the grand-average of potentials recorded by each electrode. The red lines indicate the recordings by the electrode underneath the coil (C3). The vertical dash indicates the point of time when TMS is applied. A) Responses to the stimulation of M1. B) Responses to the stimulation of shoulder. C) The upper and middle topoplots depict voltage distributions across the scalp for each peak of interest, in response to the real and control conditions, respectively. The lower topoplots illustrate the results of the cluster-based permutation tests comparing the voltage distribution of the two responses at each peak. Clusters were defined as at least two neighbouring electrodes exceeding the threshold of p-value $< 0.05$ at each point of time. Monte Carlo p-values were calculated on 5000 iterations with a critical $\alpha$ level set at $p<0.025$. The channels highlighted by blue dots belong to the clusters that showed statistically stronger responses to the real TMS condition ($p<0.025$). Three negative and two positive significant clusters were found.

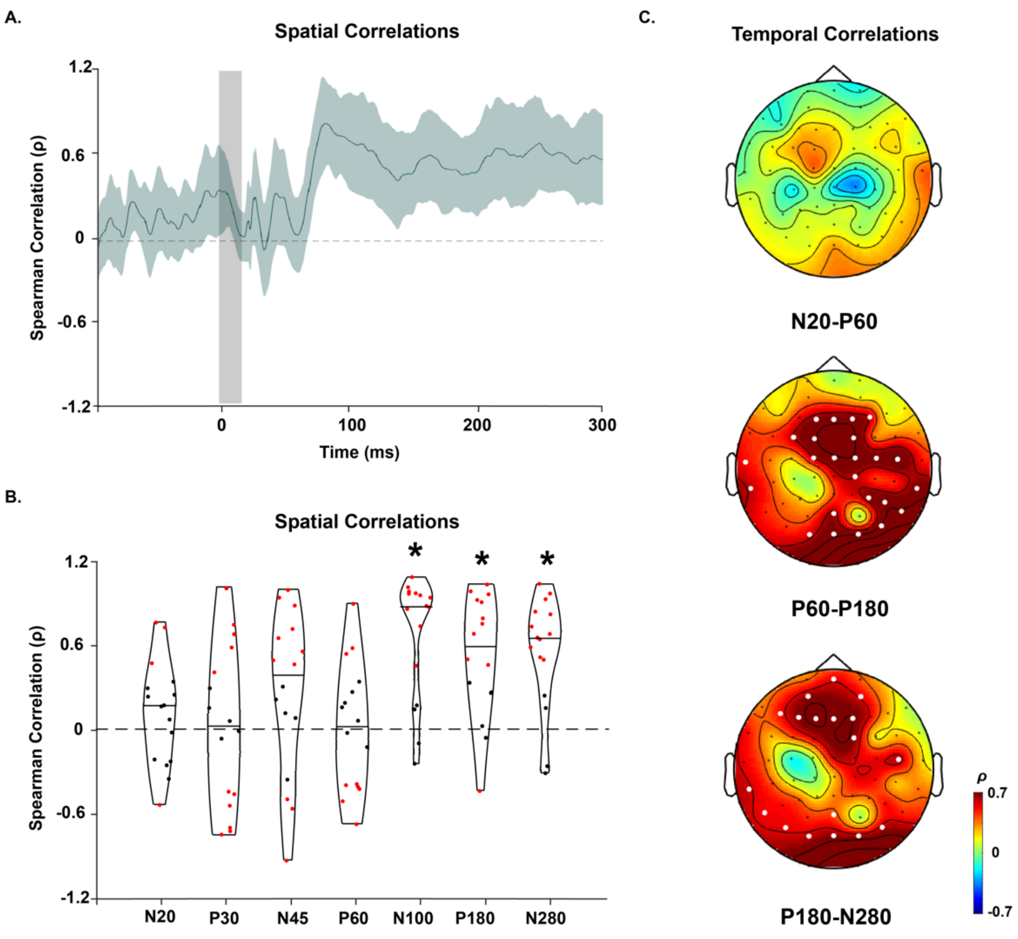


**Figure S3: Spatiotemporal correlations of TEPs and PEPs evoked by suprathreshold, monophasic TMS.** A) The spatial correlations of the potentials at each point of time from 100 ms before to 300 ms following stimulations. The green shaded area represents the 95% CIs. The vertical grey bar shows the window of interpolated potentials around stimulus. B) The distribution of spatial correlations across individuals. The dots within the violin plots represent the correlation values at for each individual. The red dots show significant positive and negative correlations respectively ($p<0.05$) and the black dots represent non-significant correlations. * indicates that correlation values differed from 0 at the group level (one-sample t-test, $p<0.05$). C) The temporal correlations of the potentials at each window of time. White dots indicate the electrodes with significant positive correlations ($p<0.05$). No significant negative correlation was found.

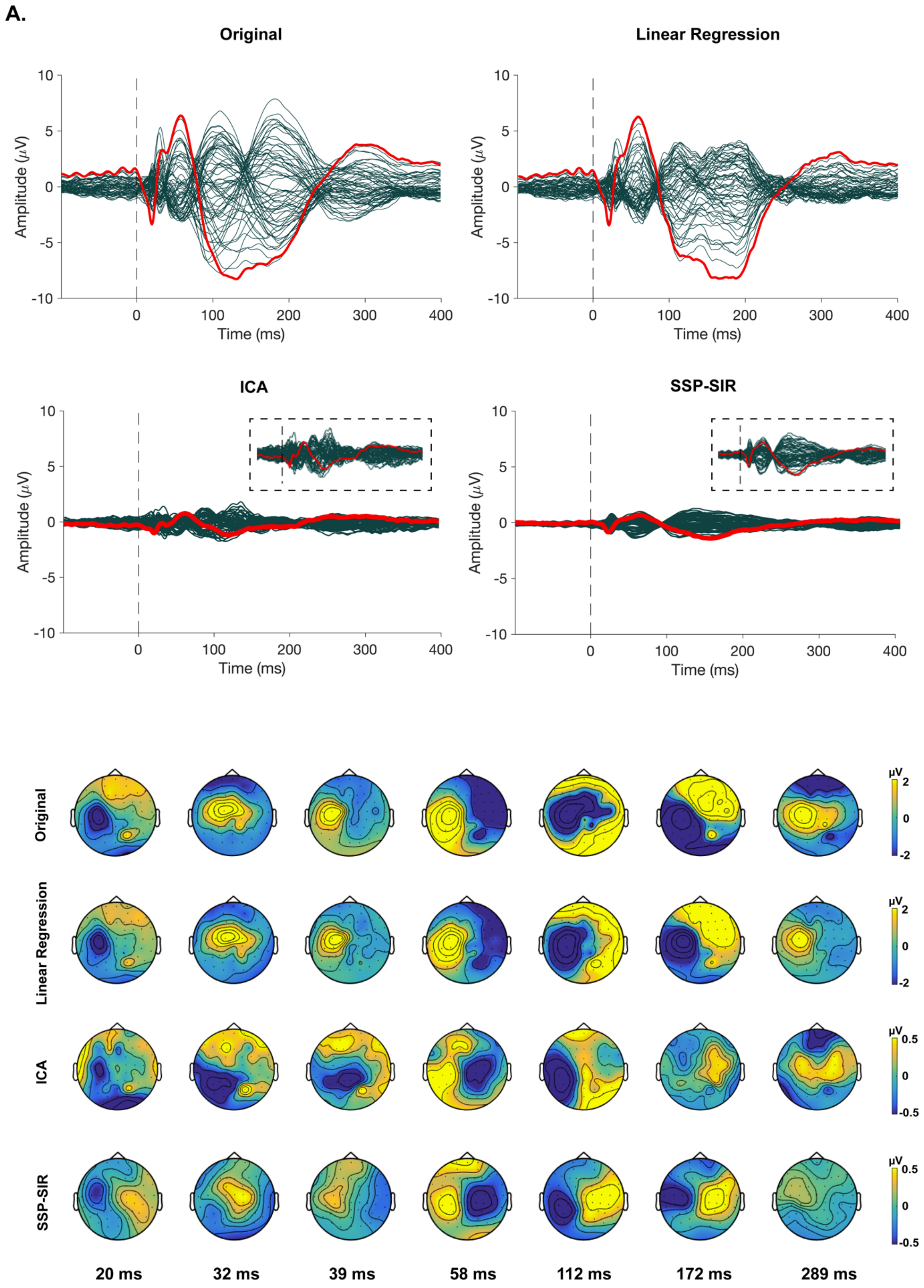
A.
Original
Linear Regression
ICA
SSP-SIR
Amplitude (μV)
Time (ms)
Original
Linear Regression
ICA
SSP-SIR
μV
20 ms
32 ms
39 ms
58 ms
112 ms
172 ms
289 ms

**Figure S4: The alterations in the spatiotemporal distributions of TEPs induced by suprathreshold and monophasic TMS before and after removing PEPs using three different filtering methods.** A) The butterfly plots demonstrate the grand-average of the potentials recorded by each electrode before (original) and after employing each filtering method. The red lines indicate the recordings by the electrode underneath the coil (C3). The vertical dash line indicates the point of time when TMS is applied. It should be noted that the y-axes are rescaled following ICA and SSP-SIR to indicate the patterns of the small TEPs more clearly. Figure insets display a magnified view of the butterfly plots used to show the patterns in the small signals (i.e. ICA and SSP-SIR) more clearly (Y axis scale is set to [-2, 2]). B). The topoplots depict voltage distributions across the scalp for each peak of interest before (original) and after applying each filter. The colour bars have been re-scaled following ICA and SSP-SIR to indicate the spatial distribution of the small TEPs more clearly.

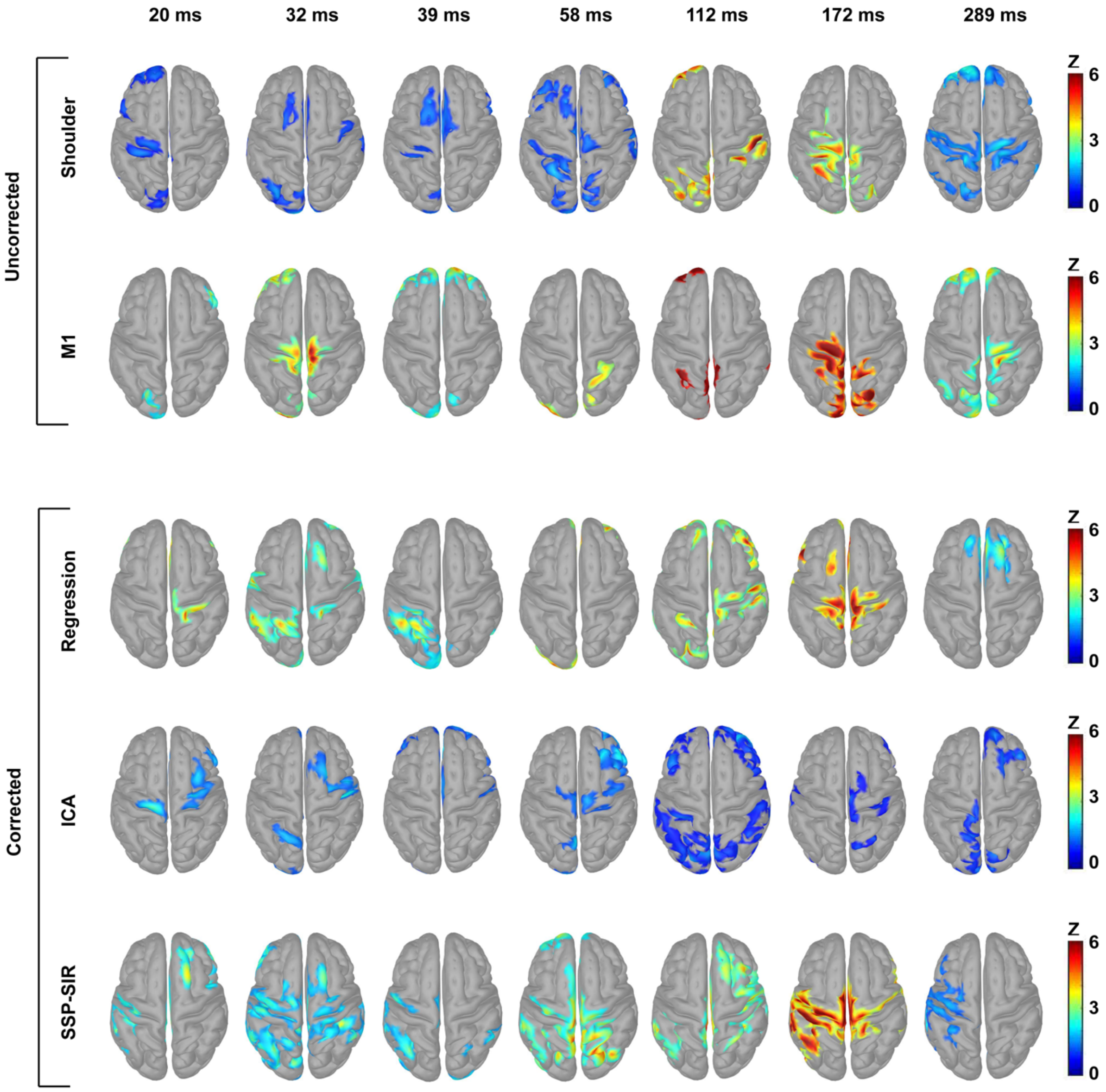


**Figure S5: The estimated source distributions obtained by applying MNE to the TEPs (evoked by suprathreshold and monophasic TMS) from different filtering methods.** MNE maps are thresholded at 40% of the maximum activity at each point of time and the minimum size for the activated regions is set to 50 vertices.

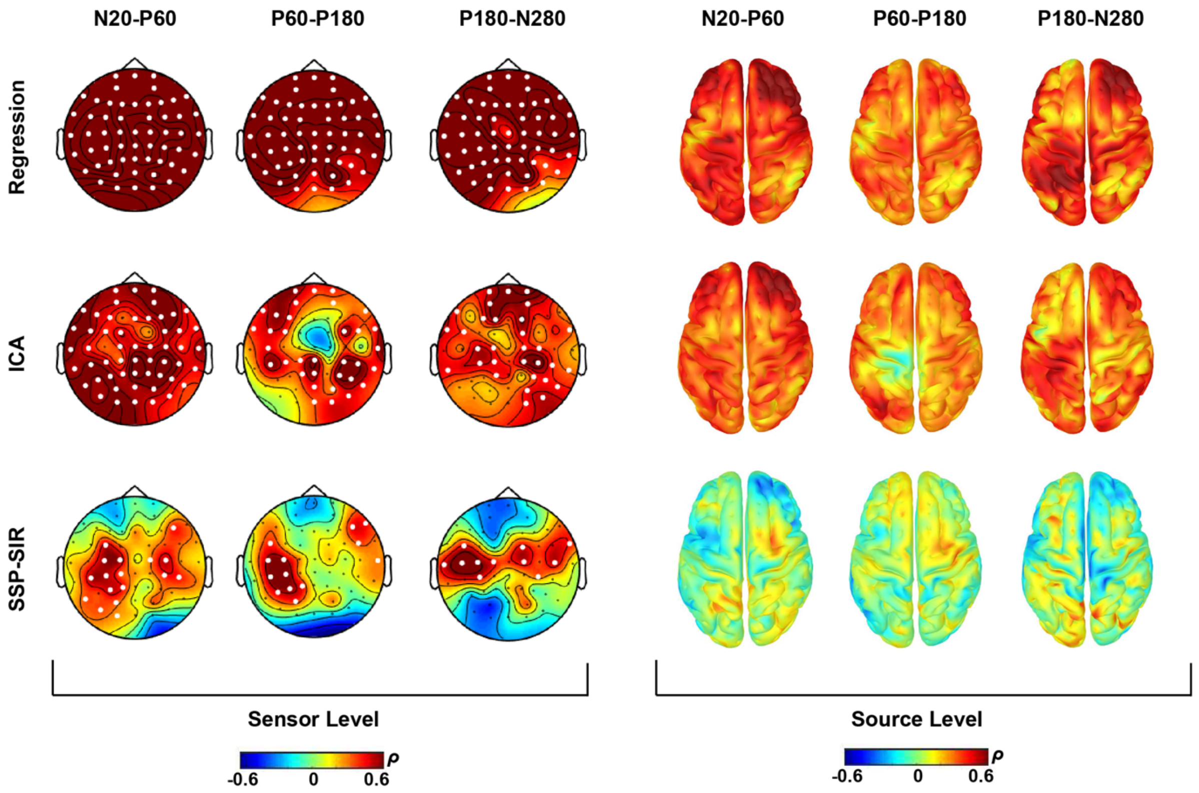


**Figure S6: Spearman correlation measures between the original and filtered TEPs (evoked by suprathreshold and monophasic TMS) at both sensor and source levels, at three different intervals.** The maps show the average of the correlation values at individualized time windows (i.e. N20-P60, P60-P180, P180-N280). A) The correlations between the original and filtered potentials recorded by each channel at each window of time. White dots indicate the electrodes with significant positive correlations ($p<0.05$). B) The distribution of the correlations between the estimated source activities at each vertex. Linear regression filtered data showed significant correlations with the original signal across the whole time and space domains. ICA, also, showed strong widespread correlations across time at both source and sensor levels. SSP-SIR resulted in substantially lower correlations (high suppression) especially around the fronto-central regions, which had shown sensory high contamination, but caused minimum suppression to the recordings around the site of stimulation especially noticeable at the scalp level.

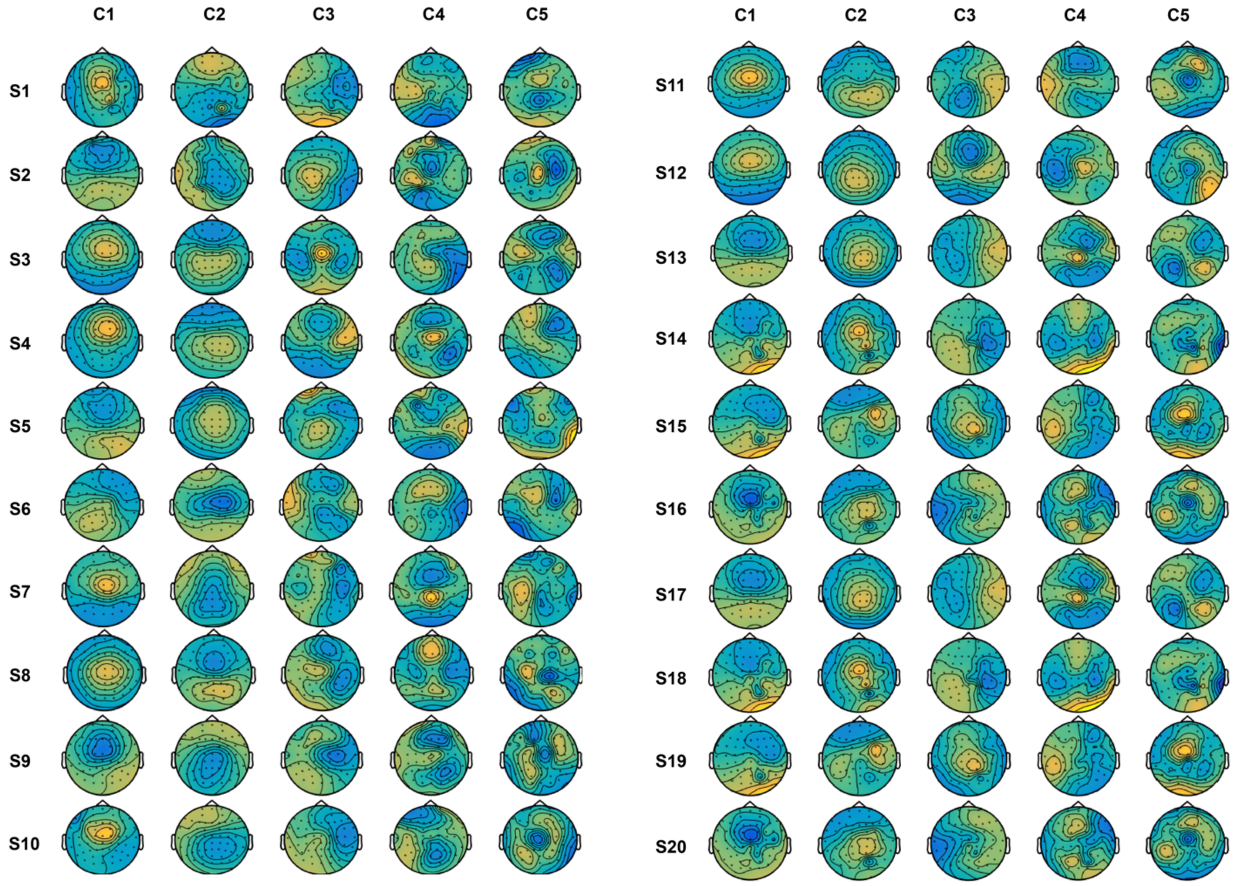


**Figure S7: Topography of the first five artifactual components (C) rejected by SSP-SIR, for each subject (S) at biphasic and suprathreshold stimulation condition.**

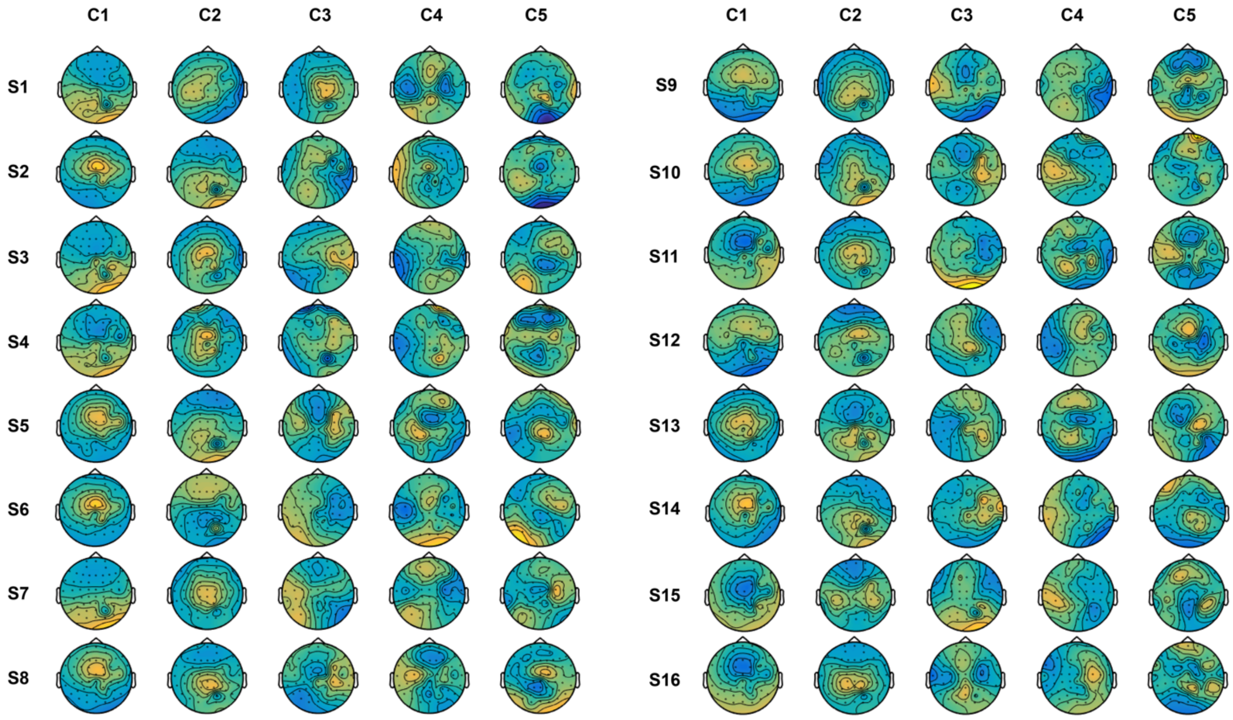


**Figure S8: Topography of the first five artifactual components (C) rejected by SSP-SIR, for each subject (S) at monophasic and suprathreshold stimulation condition.**
